# Bioengineering Fascicle-like Skeletal Muscle Bioactuators via Pluronic-Assisted Co-axial 3D Bioprinting

**DOI:** 10.1101/2024.09.06.611597

**Authors:** Judith Fuentes, Rafael Mestre, Maria Guix, Ibtissam Ghailan, Noelia Ruiz-González, Tania Patiño, Samuel Sánchez

**Affiliations:** Institute for Bioengineering of Catalonia (IBEC), Barcelona Institute of Science and Technology (BIST), Baldiri-Reixac 10-12, 08028 Barcelona, Spain; School of Electronics and Computer Science, University of Southampton, Highfield, SO17 1BJ, UK; Department of Materials Science and Physical Chemistry, Institute of Theoretical and Computational Chemistry, University of Barcelona, 08028, Barcelona, Spain; ICFO - Institut de Ciències Fotòniques, The Barcelona Institute of Science and Technology, Biomedical optics, C. F. Gauss, 3, Castelldefels, Barcelona, Spain; Biomedical Engineering Department, Institute for Complex Molecular Systems. Technische Universiteit Eindhoven, Het Kranenveld 14, 5612 AZ Eindhoven The Netherlands; Institució Catalana de Recerca i Estudis Avançats (ICREA), Passeig de Lluís Companys 23, 08010 Barcelona, Spain

**Keywords:** bioengineering, 3D bioprinting, muscle-based actuators, pluronic, co-axial printing

## Abstract

Advances in 3D bioprinting have opened new possibilities in the development of bioengineered muscle models that mimic the structure and functionality of native tissues. The combination of skeletal muscle tissue and artificial elements promotes diverse innovative solutions of interest in both the biomedical field and the development of biohybrid actuators. However, current bioengineering approaches do not fully recreate the complex fascicle-like hierarchical organization of skeletal muscle, impacting on the muscle maturation process due to a lack of oxygen and nutrients supply in the scaffold inner regions. Here we explored co-axial 3D bioprinting as a strategy towards overcoming this challenge, creating individual/non-fused filaments with controlled thickness that present a fascicle-like organization. Compared to conventional 3D-bioprinting, where cell-laden bioink is disposed by a single syringe, our Pluronic-assisted co-axial 3D-bioprinting system (PACA-3D) creates a physical confinement of the bioink during the extrusion process, effectively obtaining thin and independent printed fibers with controlled shape. Fabrication of skeletal muscle-based actuators with PACA-3D resulted in improved cell differentiation, obtaining stronger bioactuators with increased force output when compared to bioactuators fabricated by conventional 3D bioprinting. The versatility of our technology has been demonstrated using different biomaterials, showing its potential to develop more complex biohybrid tissue-based architectures with improved functionality.

## 1. Introduction

The 3D bioprinting technique allows the automated and fast-prototyping printing of cell-laden hydrogels with biomimetic designs resembling the 3D environment of native tissue. [1,2] Opposed to two-dimensional cultures, 3D bioprinting provides three-dimensional environments that better recreate the in vivo conditions in biological models, leading to an improved cell-cell interaction, division and morphology.[3] Pneumatic-extrusion-based bioprinting is the most widely used in biofabrication, exploring different biocompatible hydrogels that allow optimal cell distribution and interaction during the maturation process. This technique is based on the extrusion of a cell-laden hydrogel through a nozzle when controlled pressure is applied, using materials with enough pseudoplastic or shear-thinning behavior, which allows printing at lower pressures while protecting cells from high shear stress conditions.[4] Materials like alginate, gelatin, fibrinogen, hyaluronic acid, chitosan, poly(ethylene glycol) diacrylate (PEGDA), collagen, nanocellulose, decellularized ECM, or chemical modifications of these (i.e., gelatin methacrylate, GelMA) have been widely used alone or in combination with one another for their shear-thinning properties, stiffness modulation or biocompatibility.[2,3,5–7] More complex structures can be achieved by bioprinting multiple materials in parallel, using for example polycaprolactone,[8] polydimethyilsiloxane (PDMS)[9–11] or pluronic acid [12,13]. This last, is commonly used as a sacrificial material for obtaining overhanging structures or create vascular channels/complex 3D structures, as it can be easily washed away with cold water or PBS. Other interesting examples in pneumatic-extrusion-based 3D bioprinting have explored the use of cartilage,[14] neural tissue,[15] cardiac,[16] skin,[17] as well as multiple cell bioprinting for myotendinous tissues.[18]

Native skeletal muscle tissue presents an inherently three-dimensional architecture composed by bundles of fibers hierarchically organized in fascicles. Therefore, any tissue model fabricated that aims at recreating its complexity must consider this three-dimensional configuration, either by following 3D-bioprinting techniques[19,20] or mold-casting[21]. One of the main challenges remains on overcoming the lack of oxygen and nutrient diffusion when generating thick cell constructs. Indeed, the oxygen diffusion limit within a tissue is around 200 μm,[22] leading to a great loss of output force due to the lower myotubes density in the muscle bundle.[23] Moreover, the fascicle-like structure of native muscle has not been completely mimicked yet with 3D bioprinting due to the filaments fusion during the printing process when extruded subsequently (i.e., layer by layer). Although some examples of fascicle-like constructs generated by using of sacrificial molds and microfabrication techniques have been demonstrated,[24–26] these techniques lack the versatility and automation of 3D bioprinting methods. Recent advances in 3D printing of skeletal muscle constructs for tissue restoration have focused on innovative photo-crosslinking techniques, enabling in situ crosslinking of bioinks during extrusion.[19,20,27] This approach allows precise control over the width and shape of the printed filaments, which is crucial for the further development of fascicle-like structures. Despite these benefits, its versatility is constrained by the requirement for photo-crosslinkable bioinks. Co-axial printing, another approach commonly used for the fabrication of tubular structures and vascular scaffolds,[28–30] has also been demonstrated to allow the creation of fascicle-like tissues.[31] Lee et. al replicated muscle fascicles and the perimysium surrounding them by 3D printing a core-shell structure consisting of a photo-crosslinkable polymer mixture encapsulating C2C12 cell aggregates suspended in media. However, co-axial printing holds potential beyond the fabrication of vascular-like and core-shell structures - it can also be applied to fabricate individual non-hollow filaments of defined widths through the co-extrusion of sacrificial materials in the outer layers, generating fascicle-like structures with greater control over filament size and so homogeneity.

Specifically, in this work we propose a Pluronic-Assisted Co-Axial 3D bioprinting technique (PACA-3D) as a novel and universal method to obtain non-hollow individual filaments. This technique: i) produces homogenous filaments with controlled width down to 190 μm, ii) generates individual filaments that do not fuse with each other, creating a fascicle-like structure that improves the biomimicry of the native skeletal muscle tissue; and iii) is a flexible and universal bioprinting method, as it can potentially include different types of hydrogels and different crosslinking strategies, even collagen (one of the main components of the skeletal muscle ECM). Furthermore, we tested their functionality as a bioactuator by evaluating its force output at different maturation stages. We observed that the bioactuators fabricated by PACA-3D with a fascicle-like structure exerted an enhanced contractile behavior compared to muscle actuators fabricated with conventional 3D bioprinting. Therefore, such bioengineering tissue can be used to study muscle development, maturation or healing, [32–34] and as drug testing platform for biomedical or cosmetic purposes, [11,35] as well as being of great interest to develop more complex untethered actuators and bio-hybrid soft-robots. [36–38]

## 2. Results and discussion

### 2.1 The pluronic-assisted co-axial 3D bioprinting technique (PACA-3D)

Skeletal muscle is composed of bundles of muscle fibers (figure 1(A)), which are the basic contractile unit of the muscle and are created after fusion of myoblasts, their precursor cells, during the differentiation process.[39] Inside each muscle fiber, groups of internal protein structures, called the myofibrils, are formed.[40]. Moreover, in native tissue each bundle of myotubes is further organized in fascia or fascicles surrounded by the perimysium, a layer of connective tissue (mainly collagen) that stabilizes and separates the bundles of muscle fibers, containing from 10 to 100 of them.[41] Here we aimed to replicate this complex and hierarchical structure using PACA-3D.

**Figure 1.**
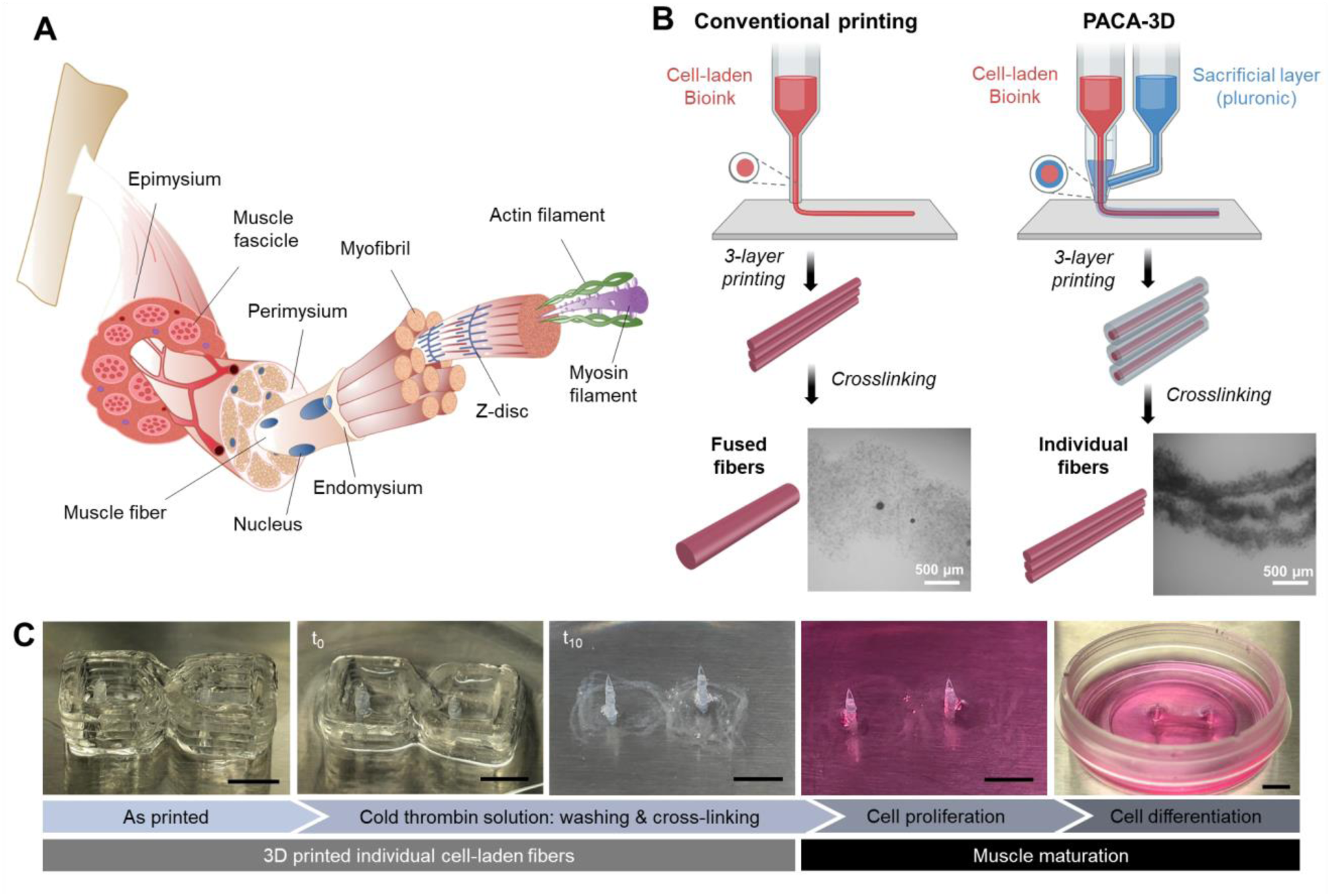
Fabrication of bundle-like structures using pluronic-assisted co-axial 3D bioprinting technique (PACA-3D). (A) 3D structure of the skeletal muscle tissue hierarchically organized in fascicles, containing several bundles of multinucleated myofibers. (B) Schematics of conventional and pluronic-assisted coaxial 3D bioprinting. In conventional bioprinting, the layers fuse together during the crosslinking process, resulting in a single filament. In coaxial bioprinting, bioink layers do not fuse due to the co-extrusion with pluronic acid (sacrificial material). Therefore, after bioink crosslinking and washing of the pluronic, thinner individual fibers are obtained. C) The printed construct is washed and crosslinked simultaneously to finally obtain individual thin filaments (images at time 0 min and 10 min, respectively). Later on, printed tissue is maintained in GM for two days and then changed to DM to induce differentiation into myotubes in the two-post system. Scale bar: 5 mm. This figure has been created with BioRender.com

Co-axial 3D bioprinting is a method that allows printing cell-laden hydrogels surrounded by a shell of another hydrogel, usually assisting the crosslinking of the inner one. One of the most common strategies for co-axial printing is the use of alginate, which is crosslinked in situ during the extrusion process by the presence of calcium chloride in the outer needle. However, this method can be difficult to regulate, as the calcium chloride flow in solution is difficult to control (requiring the use of syringe pumps to achieve sufficiently small and controlled flows) and the obstruction of the internal nozzle often occurs due to the accumulation of crosslinked alginate.[42] Here, to overcome all these difficulties, we considered an alternative approach based on using a sacrificial material as a supporting structure during the 3D bioprinting process. Towards that aim, we chose the block co-polymer pluronic acid F-127, which has successfully been used as a sacrificial ink for structures that need infilling or three-dimensional support before crosslinking.[43] However, pluronic acid has never been implemented in a co-axial system to act as shell to finely tune the thickness of the hydrogel extruded in the inner needle.

Figure 1(B) shows a comparative schematic of the working principles of conventional 3D bioprinting and our pluronic-assisted coaxial 3D bioprinting method. For this approach, pluronic is used to provide a confinement layer during printing, maintaining the fiber structure until crosslinked. Pluronic undergoes a sol-gel transition at room temperature, going from liquid below ∼ 4 °C to a gel at room temperature, making it ideal for 3D printing and obtaining a fine control of its extrusion. Compared to conventional printing, with co-axial bioprinting, thin, individual and width-controlled filaments that do not fuse with each other can be printed. With this method, there is no need to crosslink the hydrogel during extrusion, but simple physical confinement between the cell-laden hydrogel and pluronic ensures the confinement of the hydrogel. Both strategies provide initial stability of the 3D-bioprinted fibers and allow for a second-level and final crosslinking by any available approach, namely: i) UV-crosslinking; ii) temperature-assisted crosslinking; iii) enzymatic crosslinking; or iv) ionic crosslinking. Once the cell-laden hydrogel is crosslinked, pluronic can be removed by the addition of cold PBS, inducing its dissolution at low temperature. As an example, in figure 1(C) we show the process for a cell-laden hydrogel composed of gelatin and fibrinogen with C2C12 cells. The physical contact between gelatin and pluronic in their gel form helps the fibers retain their shape and not fuse with each other until crosslinking. Crosslinking of fibrinogen-based hydrogels is triggered by the addition of thrombin, which is already mixed in the cold PBS solution, both dissolving the pluronic outer shell and crosslinking the fibrinogen within the cell-laden hydrogel at once. As shown in figure 1(C), the muscle constructs are directly bioprinted and maintained in a two-posts system (flexible PDMS-based pillars) during the muscle maturation process. This setup, which has been previously reported,[10] not only provides mechanical stability and constant tension to the printed muscle, which contributes to an improved cell differentiation and muscular fiber alignment, but also serves as a platform to measure the contractile force.

The printing fidelity of a hydrogel is highly dependent on the type of nozzle used and on its gauge. Conical nozzles, usually fabricated in plastic, provide better stress profiles for 3D bioprinting since the point of maximum stress is only found at the end of the tip. On the contrary, cylindrical nozzles made of stainless steel could be potentially harmful for cells, as the shear stress increases significantly when they are too long. Previous reports on the literature of co-axial or core-shell printing have used cylindrical needles, as they are easier to assemble.[42,44,45] Here, however, we looked for a strategy that could include conical needles in the co-axial design, posing less damage for cells (detailed information about the optimization of the co-axial needles can be found in SI, figure S1).

### 2.2 The confinement methods and materials

Using pluronic acid as sacrificial material offers high versatility to implement several crosslinking strategies, not restricting its use to skeletal muscle cells, but potentially amplifying its use for many kinds of tissues that might require different hydrogel compositions. Based on these properties, two different hydrogel confinement strategies were envisaged: i) a chemical method, based on the in-situ crosslinking of alginate inside the hydrogel with CaCl2 present in the pluronic solution; and ii) a physical method, based only on physical repulsion by gelatin inside the hydrogel and pluronic outside of it, as previously mentioned (figure 2(A)).

**Figure 2.**
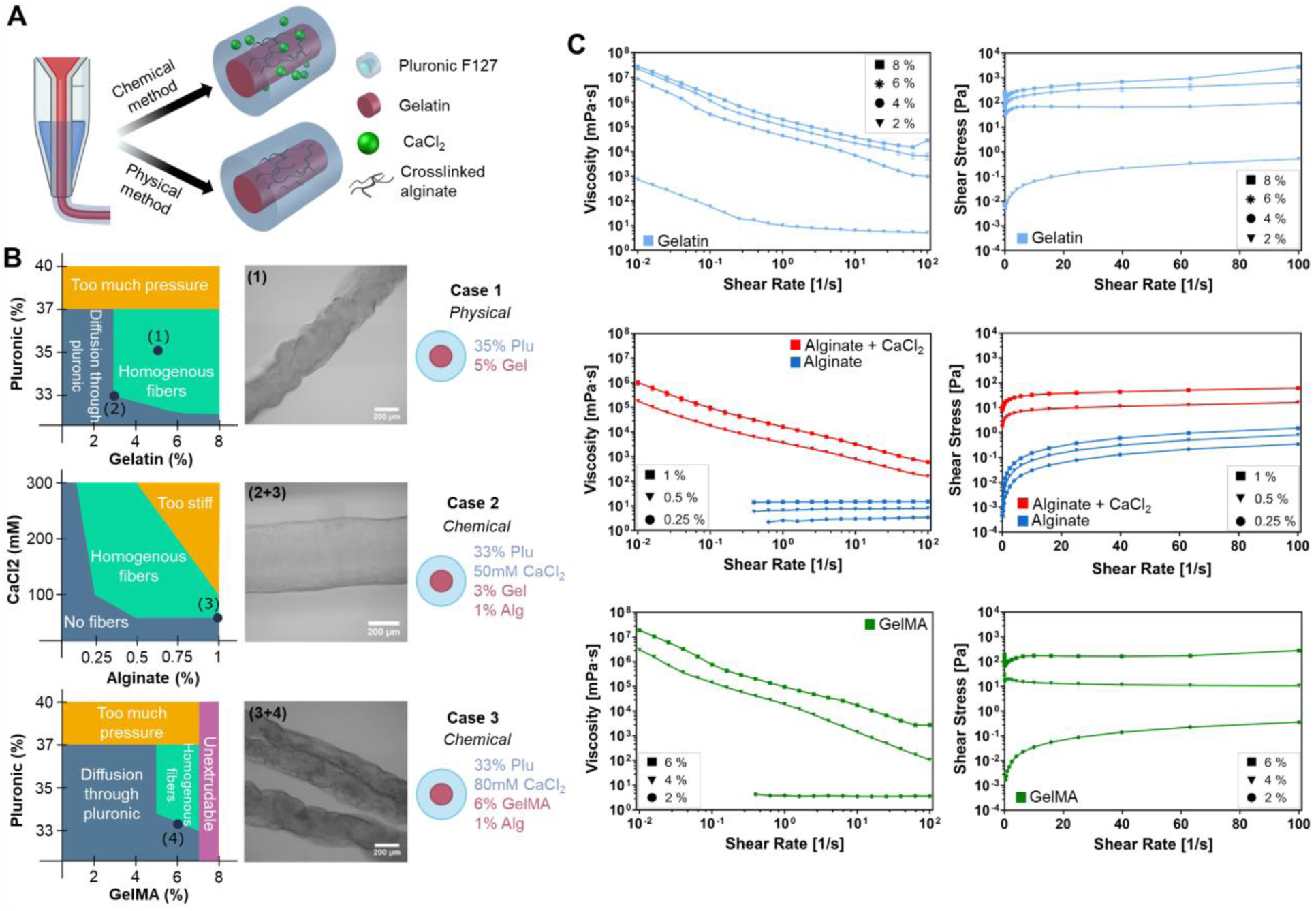
PACA-3D printing and rheological characterization of different biomaterials commonly used tissue bioengineering. (A) Schematic representation of the two different confinement methods (chemical and physical) used for co-axial bioprinting. (B) Ranges of application of different materials according to their confinement method. (C) Rheological characterization (viscosity curves) of materials at different concentrations commonly used in tissue engineering. This figure has been created with BioRender.com.

Both the chemical and physical confinement methods allow for a great degree of flexibility with the materials of choice. The chemical confinement method is ideal for materials that require ionic crosslinking, such as alginate, as these ions can be dissolved at the desired molarity in pluronic at low temperature, whereas the uncrosslinked material is added as part of the cell-laden hydrogel. Alginate and calcium chloride, one of the most widely used pairs, are a great choice for this method, but in principle any other similar combination of materials could be used (i.e., gellam gum, chitosan). For instance, in our implementation, Ca^+2^ ions from pluronic get in contact with alginate at the borders of the fiber, chemically crosslinking it to achieve a homogeneous fiber. The physical method, on the other hand, only requires the cell-laden hydrogel to be in a gel form at the bioprinting temperature. Gelatin, as we already showed, is an adequate choice for this method, as it is cheap, biocompatible, and widely used in 3D bioprinting due to its shear-thinning behavior and because it can be washed away during incubation. As with the chemical confinement method, any other material of similar gel properties could be used instead (i.e., GelMA). However, gelatin is particularly advantageous, as it can be mixed with other hydrogels that are more liquid, such as fibrinogen, collagen, Matrigel®, or alginate, serving as supporting hydrogel to obtain very thin fibers, something difficult to achieve with low viscosity hydrogels.

The versatility to adapt PACA-3D to different confinement schemes and crosslinking methods leads to a great variety of combinations of materials depending on the specific requirements of cell lines and potentially covering a wide area of applications. Table S1 shows a qualitative overview of some of the mostly used materials in 3D bioprinting that can be implemented in both of our strategies, like GelMA, fibrinogen, and collagen or Matrigel®. In the corresponding discussion in the Supplementary Information, we discuss the best approaches when combining these materials to obtain high-quality individual and thin filaments with the co-axial methods, as well as their pitfalls, caveats, and opportunities. The summary presented in table S1 is a fast guide to understand the possible material combinations according to their confinement method, but the reality is much more complex. The relative concentrations of the confinement materials, as well as that of pluronic and even CaCl_2_, will greatly influence the applicability of the co-axial printing method, as depicted in figure 2B. Regarding the physical confinement method, the concentration of gelatin with respect to the concentration of pluronic needs to be adjusted to avoid bio-ink diffusion in the outer shell. In general, gelatin at concentrations below 3% (wt/v) is liquid enough to diffuse through pluronic. At the same time, pluronic at concentrations below 33% (wt/v) is not viscous enough to prevent bio-ink diffusion. Therefore, the ideal range of concentrations for gelatin to obtain homogeneous fibers, as the one shown in case 1, is between 3-8% (wt/v); for pluronic, it is between 33-37% (wt/v). Higher concentrations of both pluronic and gelatin can still be successfully used. However, they might require a pressure range too high for 3D bioprinters (depends on the specific bioprinter model) and cells might suffer too much shear stress during extrusion. The chemically assisted method, based on alginate crosslinking on extrusion, provides the improved fibers homogeneity compared to physical confinement, as shown in figure 2(B). Moreover, it does not present clogging problems since the flow of pluronic can be carefully controlled by the applied pressure. Alginate concentrations ranging from as low as 0.25% (wt/v) can yield to homogeneous fibers if the concentration of CaCl2 is high enough and other materials that increase the viscosity are present in the mixture. Higher concentrations of alginate, even surpassing 1% (wt/v) can, naturally, produce very homogeneous fibers, but its stiffness might be too high for the cells, considering that other biocompatible polymer with attachment sites, like fibrinogen or collagen, should be included in the mixture. If needed, alginate can be removed after crosslinking of the main biomaterial by the addition of ethylenediaminetetraacetic acid (EDTA) for 5 min. [44,46]

Finally, another possible strategy consists of a hybrid chemical-physical confinement method that uses both gelatin and alginate at smaller quantities. Case 2 in figure 2(B) shows one of those examples, where alginate at 1% (wt/v) was ionically crosslinked with 50 mM of CaCl2 (point 3), with gelatin at 3% (wt/v) as extra support (point 2). Notice how the fiber is much more homogeneous than case 1, in which 5% (wt/v) of gelatin was used (point 1). Therefore, decreasing the gelatin concentration together with the addition of alginate gave to the fiber it a smoother shape. GelMA can also be used to achieve physical confinement, as it also presents gel properties. However, gelatin and GelMA gelification are slightly different due to the methacrylation process taking place during GelMA synthesis.[47] This can be seen by the relationship between pluronic and GelMA concentrations. GelMA gelates at higher concentrations than gelatin, requiring a minimum concentration of 5% (wt/v), compared to the minimum of 3% (wt/v) for gelatin (figure 2(B)). Regarding GelMA, the optimal concentration range to obtain homogeneous fibers is 5 to 7% (wt/v). For concentrations higher than this, the mixture is completely unextrudable. It should be noted, that in case of gelatin the printable concentration range is from 3 to 8% (wt/v) being broader than GelMA. Also, homogeneous fibers can be obtained when adding alginate to the mixture, as shown in case 3 (point 3 and 4).

Figure 2(C) shows the rheological properties of hydrogels made of gelatin, GelMA and alginate at different concentrations. Viscosity curves of gelatin and GelMA indicate the non-Newtonian behavior of these materials at a concentration above the 2% (wt/v), as shown by the decrease of viscosity when higher shear rates are applied. Indeed, this behavior corresponds to shear thinning materials, which are the ones commonly used in extrusion-based 3D printing for allowing the printing of homogeneous filaments. Interestingly, both gelatin and GelMA at 2% (wt/v), although GelMA specially, do not present non-Newtonian properties, as the viscosity remains mostly constant when increasing the share rate, meaning that at low concentrations the hydrogel is mostly liquid and therefore not suitable for extrusion printing. Something similar is observed with alginate. Alginate at the concentrations reported in the hereby investigation shows Newtonian properties (i.e., viscosity is constant regardless the shear rate), but when calcium chloride is added to the solution inducing the alginate crosslinking, the final hydrogel presents a shear thinning behavior similarly to gelatin and GelMA.

### 2.3 Biofabrication and characterization of skeletal muscle bioactuators with fascicle-like structures using PACA-3D

#### 2.3.1 Characterization of 3D printed individual filaments with controlled width

The descriptions of the previous sub-section are of a universal character; this PACA-3D system, together with any of the confinement methods, or their hybrid combination, can be used to create thin and homogeneous filaments of controlled width. Moreover, a wide variety of biomaterials, such as fibrinogen, Matrigel®, collagen, or GelMA, with different crosslinking strategies, like enzymatic, ionic, UV light or temperature, can be used to provide the suitable three-dimensional environment for cell survival, depending on the specific needs of each cell type. One of the most straightforward applications of this technique is found in the 3D bioprinting of skeletal muscle cells, which can benefit the most from the fibrillar structure, inducing a self-organization in a fascicle-like manner, as in native tissue.

With any of the two confinement methods mentioned in the previous section, chemical and physical, filament width could be carefully controlled to the desired dimensions by simply modifying the pressure applied on the hydrogel. The desired width is 200 µm for presenting the oxygen diffusion limit within a tissue, being crucial to work within that width values to improve cell viability and therefore, cell differentiation and muscle maturation. Figure 3(A) shows the range of width that was available for each case. For the chemical confinement method, the control was finer than for the physical one, achieving diameters as low as the nozzle diameter (200 μm) for 60 kPa of pressure. However, no continuous filaments were obtained applying less than 60 kPa. The physical confinement method, on the other hand, could span from around 200 μm width when 50 kPa were applied to 330 μm when applying 80 kPa, meaning that thinner filaments could be obtained at any working pressure. In any case, the variability was low, as demonstrated by the error bars, indicating that the printed filaments were homogenous. Notice, however, that the exact range of pressures is highly dependent on the characteristics of the bioprinter, tubes, nozzles, and pressure pumps, as well as the hydrogel concentration. Despite the applied pressure range, our method shows great flexibility and homogeneity during bioprinting. Contrary, when using conventional 3D bioprinting, the diameter of the printed filaments was more than double in some cases than the ones obtained with coaxial printing, even at low pressures, demonstrating the effectivity of the PACA-3D printing strategy for the fabrication filaments of controlled width.

**Figure 3.**
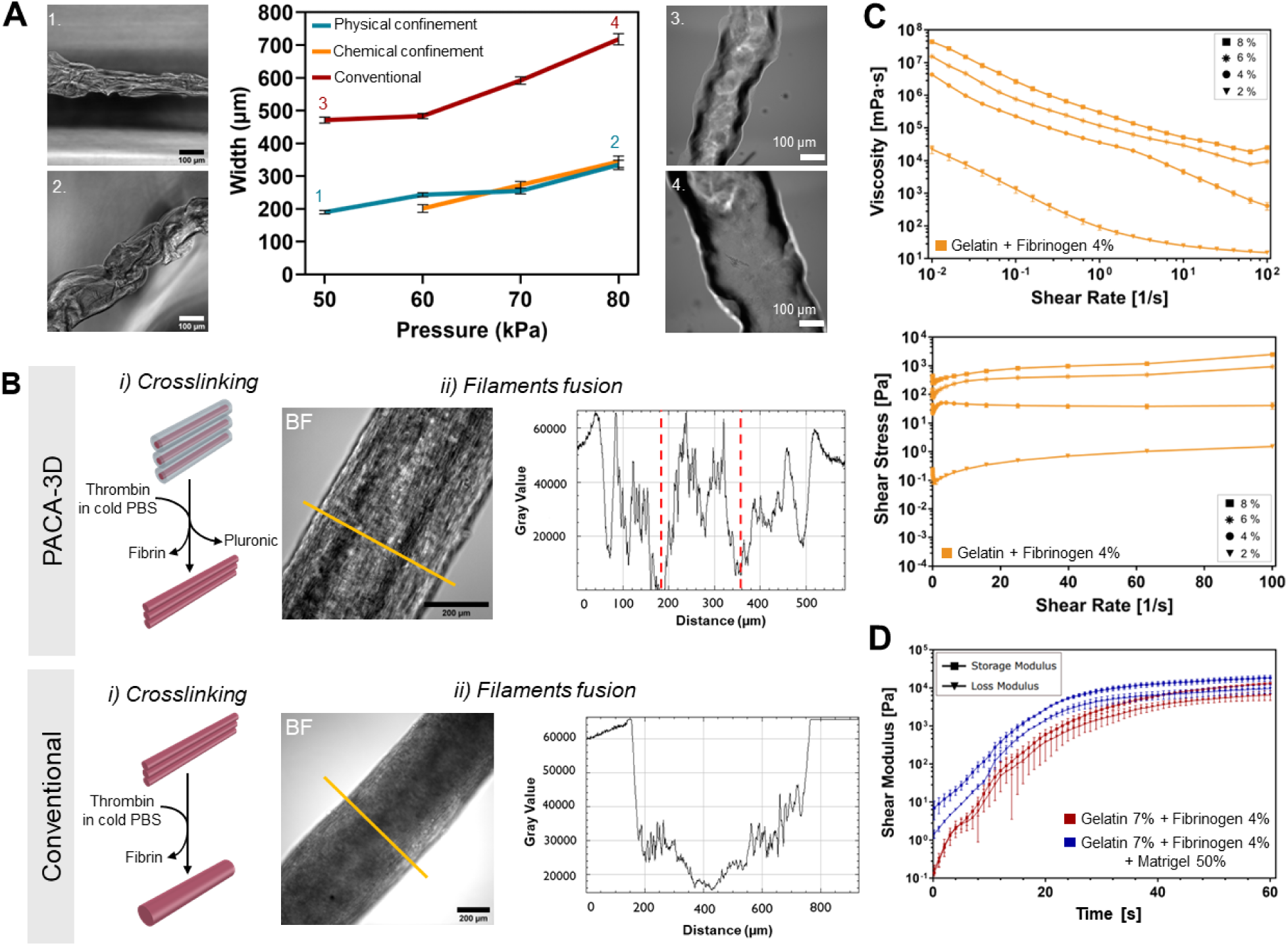
Characterization of the gelatin- and fibrinogen-based bioink for 3D bioprinting of skeletal muscle cells. (A) The filament diameter of a hydrogel composed of gelatin 7% (wt/v) and fibrinogen (4% wt/v) can be carefully controlled by modifying the extrusion pressure during 3D printing. Error bars represent standard error of the mean. (B-i) Schematic of the enzymatic crosslinking process of the cell-laden bioink composed of 5×10^6^/ml of C2C12 myoblasts, 7% (w/v) gelatin and 4% (w/v) fibrinogen with thrombin after 3D bioprinting. Regarding PACA-3D (top), the cold thrombin solution crosslinks the fibrinogen while pluronic acid is being dissolved, resulting in thin individual filaments. (B-ii) Bright Field (BF) microscopy images of a skeletal muscle bioactuator fabricated with PACA-3D (top), where 3 layers can be distinguished, and conventional bioprinting (down) where no individual filaments can be observed. On the right, the intensity projection graphs of the microscope images (yellow section)**. (**C) Viscosity curves (top) and flow curves (bottom) of gelatin-fibrinogen hydrogels at different gelatin concentrations, with fibrinogen fixed at 4% (wt/v). (D) Evolution of shear modulus of gelatin-fibrinogen hydrogels with and without Matrigel® over time, starting from the point at which fibrinogen is added to the mixture, which is kept at 37 °C to induce Matrigel® crosslinking as well.

Another key advantage of our technology is the fabrication of individual filaments which do not fuse together during crosslinking due to the physical confinement provided by pluronic acid during the extrusion process (figure 3(B-i)). This enables the fabrication of skeletal muscle tissues with fascicle-like structures that resemble native tissues, as shown in figure S4. Although a certain degree of filament fusion has taken place during the 9 days of muscle cell differentiation, both for PACA-3D and conventional printed filaments (figure 3(B-ii)), it is important to note that the key difference is that in conventional bioprinting such fusion event occurs during the crosslinking process, leading to a homogeneous fusion within the filament that results in one single construct. For PACA-3D printing fusion occurs later, during the cell differentiation stage, most likely due to the ECM remodeling done by healthy cells. Indeed, a closer examination when evaluating the intensity profile on images acquired by both fabrication techniques shows significant differences between filaments. The intensity profile of the PACA-3D printed filaments shows hills and valleys, consistent with a non-homogeneous hydrogel density, whereas the conventionally printed filaments show a single valley, indicating a single fused filament with homogeneous crosslinking. We hypothesize that, despite this later fusion, individually printed filaments still retain some of their individuality with points of lower density, potentially improving nutrient diffusion to the cells. Certainly, PACA-3D shows greater viability compared to conventional bioprinting, as depicted in figure S1(C) for 3D bioprinting C2C12 myoblasts in a co-axial system with and without pluronic as a confinement material. After 24 h, the percentage of viable cells in the co-axial system using pluronic has been slightly reduced to 80%; however, without using pluronic the viability is much lower after the same time, reaching almost 50%.

#### • **2.3.2.** Characterization of the printability of bioinks for skeletal muscle fabrication

For skeletal muscle tissue, fibrinogen has extensively been used, either by itself or in combination with others, like Matrigel®, collagen and/or gelatin.[3,48] Supplementary figure S3 shows the range of application of some of these mixtures, focusing on their concentrations as function of the attachment sites that they provide and the density of the matrix. The relative combination of these materials needs to be carefully considered, as the resulting bioink must be extrudable, homogeneously mixed, and after crosslinking, the matrix must not be too dense for cell proliferation. For instance, gelatin was included in all the bioinks as it provided the required viscosity for extrusion. When combining fibrinogen with gelatin, the ideal ratio should be kept between 3 and 8 % (w/v) of gelatin and 1% and 4% of fibrinogen to obtain a bioink with proper density and enough attachment sites (cases 1 and 2). If Matrigel® is mixed at a 30% (v/v) concentration with gelatin 7% (w/v), we suggest decreasing fibrinogen to a range as low as 10 mg/ml (case 3). However, if fibrinogen concentration is too low, the density of attachment sites of Matrigel® and its stiffness is not high enough for proper cell differentiation, as case 4 shows (50% (v/v) Matrigel®, gelatin 7% (w/v) and no fibrinogen). Matrigel® and gelatin present a liquid consistency at 4 °C and 37 °C, respectively. This poses a challenge when both materials are mixed, as the crosslinking temperature of Matrigel® is at 37 °C. Therefore, the mix of these two materials can lead to a non-homogeneous bioink.

Figure 3(C) shows the viscosity and flow curves of a hydrogel mix composed of 4% fibrinogen and gelatin at different concentrations, ranging from 2 to 8% (wt/v). When comparing these graphs with the ones obtained from gelatin hydrogels (figure 2(C)), no significant differences in viscosity when gelatin is at 4%, 6% or 8% (wt/v) were observed. However, at 2% (wt/v) of gelatin we observed an increase in viscosity when the fibrinogen is added to the solution, indicating that fibrinogen improves the printability of hydrogels at low gelatin concentration range. Moreover, the shear modulus of gelatin- and fibrinogen-based hydrogels with and without Matrigel® over time was evaluated. The graph in figure 3(D) shows the transition from liquid to gel state when fibrinogen is added to the mixture. The chemical crosslinking of fibrinogen with thrombin and the thermal crosslinking of Matrigel® and gelatin induces an increase of storage modulus over loss modulus overtime. When Matrigel® was added to the hydrogel mix, there was a faster transition to gel, indicated by a storage modulus higher than the loss modulus acquired during all the measurement time. This could present difficulties in terms of bioprinting, as the irreversible gelation of Matrigel® could lead to partial or complete blockage of the syringe.

These results suggest that the gelatin-fibrinogen mix is the optimal hydrogel for cell proliferation, due to its rheological behavior during the printing process and reversable gelation, both associated to gelatin presence and the abundance of cell attachment sites attributed to fibrinogen. Therefore, the combination of 7% (w/v) of gelatin and 4% (w/v) of fibrinogen was used for the 3D bioprinting of skeletal muscle tissues as described in the next section.

#### 2.3.3 Contraction force evaluation of skeletal muscle bioactuators

The functionality of the muscle tissues is evaluated in terms of contractile behavior, being assessed to understand if the creation of a more biomimetic muscle tissue with hierarchical structure will impact on the resulting force output. Figure 4(A) shows a schematic of the two-post system used as force measurements platform (i), where bending of the post due to the force contraction of the muscle is video-tracked (ii) and analyzed (iii) to obtain the post displacement graphs, where (iv) the single-twitch contractions of tissue are observed (movie S1). Then, Eurler-Bernoulli’s beam bending equation was used to correlate the displacement with the contraction force (more details in Materials and Methods section). Force measurements revealed that the muscle tissues fabricated by PACA-3D printing presented higher force output throughout the experiment timeline, especially at day 12, when 3 times stronger contraction force was obtained (figure 4(B) and movie S2). This significant increment in the force output suggests that the fabrication of hierarchically organized muscle tissue enhanced the myotubes maturation, which could be relate to an improved sarcomere remodeling during contraction.[49] The effects of several variables, such as number of printed filaments, differentiation day, printing approach or filament size, which directly affect the muscle tissue force generation were analyzed by a linear mixed effects model (figure 4(C)-S5). We performed a linear mixed-effect model analysis to determine the individual effects of different parameters in the final bioactuator output force, specifically the type of bioprinting, the differentiation day and the number of layers printed. The mixed-effect model is well suitable for this kind of experimental design as it takes into account both fixed effects (those parameters that are controlled and manipulated) and random effects (those that are not controlled but can influence the outcome, such as sample-to-sample variability), which are common when dealing with biological samples. In our experimental setup, we tested different sources of random effects (or random variability) in our samples: i) from the sample ID (each individual sample); ii) the cell passage (always 3 or 4); and iii) the printing ID (set of samples that were printed on the same day). We evaluated these three models (table S2) and, among them, the model accounting for variability in sample ID outperformed the others both in terms of explaining the between-group variance (with an Intraclass Correlation Coefficient, ICC, of 0.45) and the overall variance (with an R^2^ of 0.63). Thus, ca. 45% of the variance in the logarithm of the force output can be attributed to differences between the samples, and the model overall accounts for 63% of the variability the logarithm of the force. Therefore, we deduced that the cell passage (at least between passage 3 and 4) and the printing day were not significantly influencing the force output of the bioactuators, but sample-to-sample variability was.

**Figure 4.**
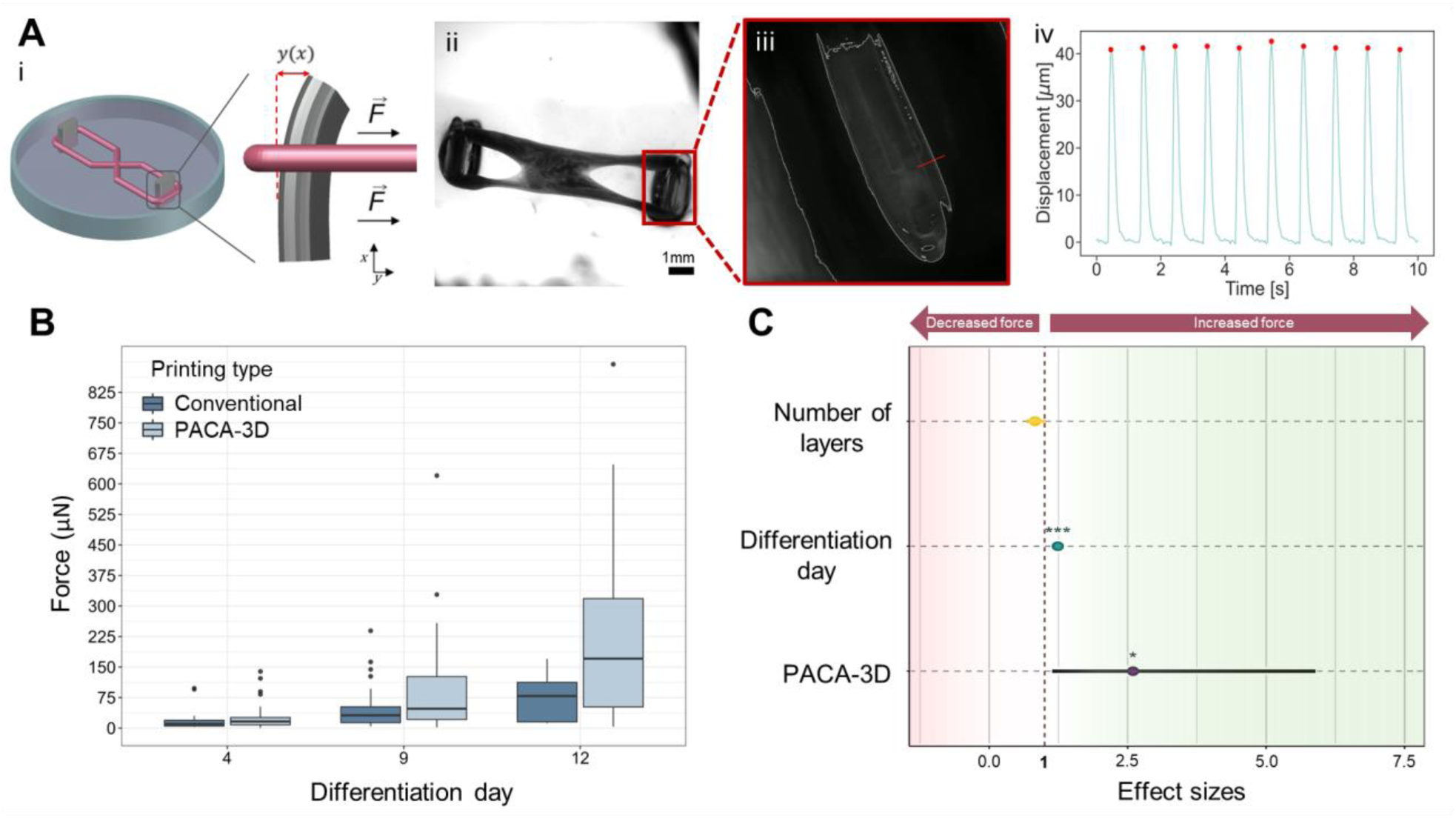
Characterization of the force contraction of the bioactuators. (A) Force monitoring mechanism schematic depicting the evaluation of force output from the 3D-bioengineered skeletal muscle actuators under electrical pulse stimulation (EPS). (B) Muscle actuator assembled on force measurement platform. Videos from the contracting muscle are taken, then analyzed via Python. Post silhouette is marked in white, and a perpendicular red line is created to determine the post displacement generating a graph A-iv) that shows the single-twitch contractions. B) Force output from muscle-based actuators fabricated using PACA-3D and conventional 3D bioprinting techniques, at day 4, 9 and 12 of differentiation. (C) Mixed effect model on the force output depending on different variables. Variables with effect sizes below 1 have negative impact on the force (red area). Contrary, effect sizes above 1 indicate that the variable presents a positive effect on the force output (green area).

Our final linear mixed model was thus described as:

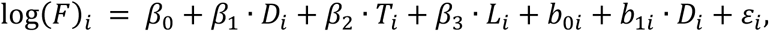

where log(*F*)_*i*_ represents the logarithm of the force for sample *i*, *²* are the model fixed effect coefficients, *D*_*i*_is the differentiation day, *T*_*i*_is the type of printing (where 0 is the conventional printing, the baseline, and 1 is coaxial prating), *L*_*i*_ is the number of layers, *b*_0*i*_and *b*_1*i*_are the random intercept and random slope for each day (respectively) model, we not only allowed for a random effect in the intercepts of the model to account for sample-to-sample variability, but also for a random slope for differentiation day, to account for the fact that different samples might follow slightly different differentiation patterns as days pass. The results of the model are shown in table S3 and table S4 and the model diagnostics in figure S6 that demonstrate the data and model fulfil the assumptions of normality and homoscedasticity.

Figure 4(C) shows the size effect (after exponentiating to improve interpretation; see table S4) of the three fixed effects: the differentiation day, number of layers and type of bioprinting. It could be expected that including more layers in the construct would lead to higher force output, however the results indicate otherwise. Printing more layers had a small, albeit negative, effect on the force, not statistically significant – for each layer the model estimates a reduction of force output by a factor of 0.83. This suggests that muscle tissues with wider diameters could still suffer from poor nutrient diffusion to the interior due to filament fusion after printing (see figure 3(B)), causing apoptosis or reduced differentiation. However, using the PACA-3D printing method compared to the conventional one produces a great improvement in the force output, nonetheless. Using the PACA-3D method resulted in an average of 2.6 times more force, statistically significant, with a large confidence interval that was skewed towards even higher values. As expected, the differentiation day also increased the force output, in this case by a factor of 1.2 per day. These results show that there is a great advantage by using the PACA-3D method to achieve more powerful muscle tissue, although there are still limitations to overcome. Although our method can prevent the fusion of the filaments during crosslinking, and thus improve the differentiation of the fibers, it cannot avoid the fusion of the filaments during differentiation, and adding more filaments into the bioactuator does not seem to increase the force output as could be expected.

#### 2.3.4 Biological characterization of skeletal muscle bioactuators

As previously mentioned, skeletal muscle tissue is constituted of a mixture of differentiated multinucleated fibers, known as myotubes or myofibers, that represent the basic contractile units and are formed by the fusion of myogenic cells called myoblasts, conforming a fascicle-like structure (figure 5(A)). We assessed the degree of tissue differentiation and maturation with both fluorescence microscopy and real-time quantitative polymerase chain reaction (RT-qPCR), as shown in figure 5(B) and (C), respectively. Immunofluorescence staining of Myosin 4, also known as Myosin Heavy Chain IIb (MyHCIIb), and cell nucleus (Hoechst) was performed on day 12 of differentiation. Myofiber size and nuclei position changes during myofiber maturation, observing a change in the nucleus distribution being at the center in early maturation stages and moving to the periphery in fully matured myofibers.[50] Highly aligned multinucleated fibers were obtained in both types of samples, demonstrating proper muscle maturation (figure 5 (B)), also reflected on the myotubes alignment, evaluated by the angle dispersion of both myotubes and nucleus after a fast Fourier transform (FFT) of a set of immunofluorescence images (complementary data in Figure S6). The nucleus of the conventionally printed tissues are significantly rounder (i.e. present higher diameter) and tend to be found closer to the center of the myofiber, which is associated with early differentiation stages. [50] Myotubes in conventionally printed samples are significantly wider, which is normally correlated with muscular hypertrophy.[51] It is unclear, however, whether the same level of hypertrophy can be obtained in lab-grown skeletal muscle tissue, especially since they do not contain satellite cells, which play a main role in in muscle regeneration and maintenance, and contribute to build up hypertrophic muscle when needed (i.e. after exercise).[52–55] Previous reports shown that the strongest tissue constructs generated in absence of satellite cells were thinner.[38,56,57] Then, it is possible that myotube and nuclei organization when being in the spreading phase are able to elongate the tissues but not grow them wider, as the repair-and-growth mechanism mediated by satellite cells cannot take place. Therefore, we hypothesize that the myofibers produced through PACA-3D printing reach a higher level of maturation compared to those generated by conventional printing, which aligns with the stronger forces observed in PACA-3D samples.

**Figure 5.**
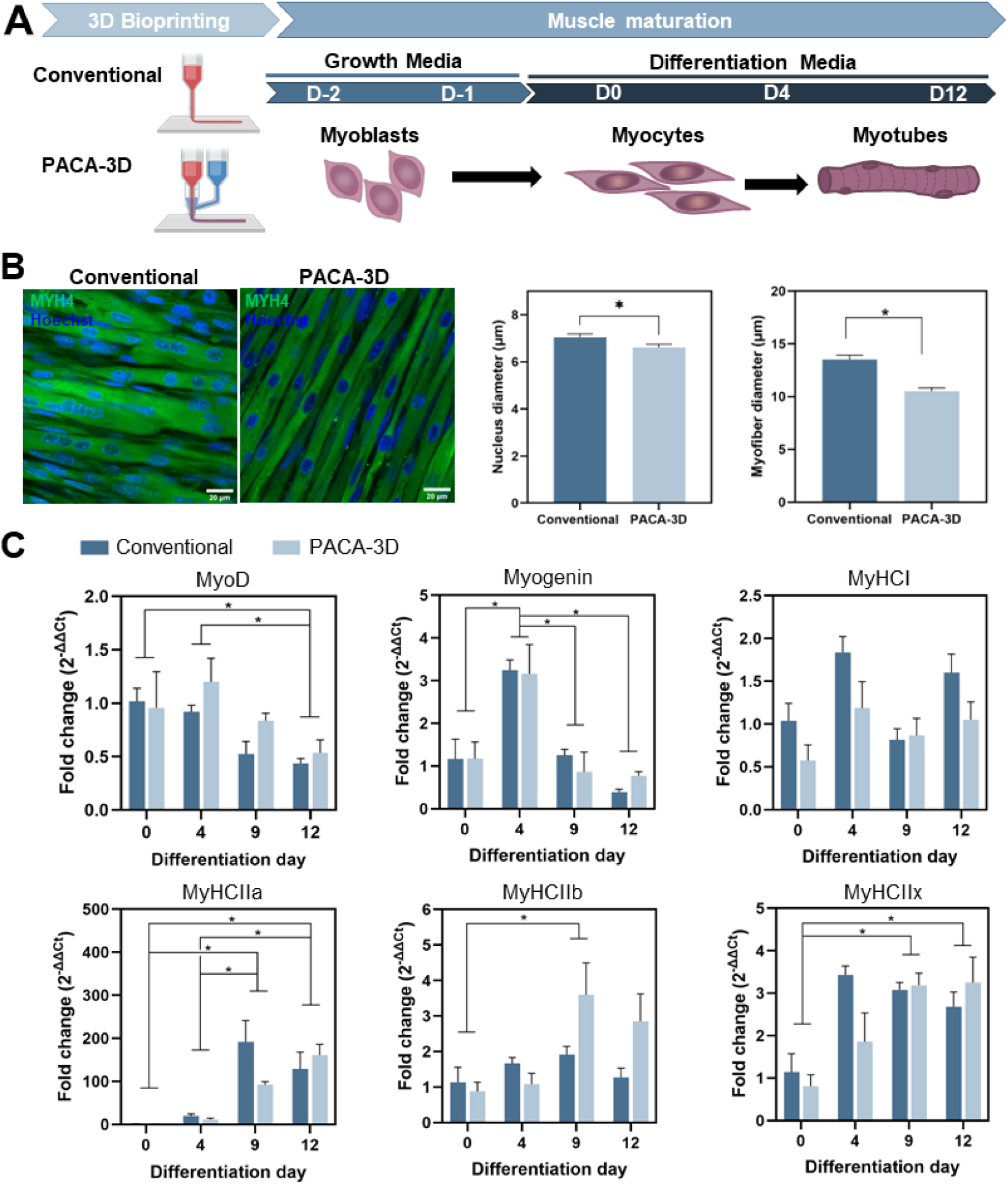
Characterization of fascicle-like cell-laden hydrogel and evaluation of myotube organization and differentiation. A) Timeline of the differentiation process of the 3D bioprinted skeletal muscle bioactuators. (B) Immunostaining of myosin 4 (MHY4, green) and Hoechst (cell nucleusn blue) after 12 days of differentiation of both conventional and coaxially printed bioactuators. Graphs display the differences in nucleus and myotube diameter between samples. (C) Gene expression analysis by RT-qPCR of the differentiation process of bioprinted muscle actuators. All values are represented as mean ± SEM. ∗p < 0.05. A non-parametric Student T-test was performed for the analysis of nucleus and myotubes diameters (N=4-6 images from a sample printed with conventional and PACA-3D) (B). A two-way analysis of variance (ANOVA)) was performed in RT-qPCR analysis (C) with N= 3 independent repeats.

The expression of myogenic markers (i.e., MyoD and Myogenin), as well as the maturation-related genes (i.e., MyHCI, MyHCIIa, MyHCIIb, and MyHCIIx), was evaluated at days 0, 4, 9 and 12 of differentiation. Generally, gene expression patterns observed in figure 4(C), correlate with our previous publications.[10,36] For instance, downregulation over time of the myoblast marker, MyoD, indicates the differentiation of these cells into myocytes and myotubes. The typical peak of expression of Myogenin, a myocyte marker gene, at day 4 is also observed. In mature skeletal muscle tissues, we can find two types of fibers: slow-twitch and the fast-twitch fibers, also known as type I and type II, respectively. MyHCI is associated to slow contraction speed (i.e., low ATP-ase activity) but provides resistance to fatigue, while MyHCIIa isoform is related to higher twitch speeds (i.e., high ATP-ase activity) that present less fatigue resistance.[58] Therefore, the functionality of skeletal muscle depends on the amount of the different fiber types found in the tissue. Interestingly, while co-axial printed tissues showed a progressive upregulation of maturation markers MyHCIIa and MyHCIIx over time, in conventionally printed samples we observed the maximum expression of MyHCIIx at day 4 and a peak of expression of MyHCIIa at day 9. These results could suggest that conventionally printed samples reached the maturation stage earlier than PACA-3D samples, although images from immunofluorescence staining of MyHCIIa suggest advanced maturation in comparison with conventional tissues. Focusing on MyHCIIb, in both coaxial and conventional printed tissues we found the highest expression levels at day 9. Notably, the expression of this gene in coaxial samples is greater than in conventional samples here and on day 12, which indicates that more fast-twitch IIb fibers are found in tissues fabricated with PACA-3D. Regarding the expression of MyHCI, we observed a downregulation at day 9 that together with the peaks of expression of MyHCIIa, MyHCIIb and MyHCIIx on the same day, indicates a slow-to-fast myosin transition in the myotubes when the maturation stage is reached.

Although there is no statistical difference between the muscle tissues fabricated with PACA-3D and conventional printing, probably due to the high variability between samples, we can conclude that the differentiation process occurs as expected, with the up regulation of MyoD and Myogenin in early stages. Regarding the maturation process, we observed that some of the type II markers are more expressed in PACA-3D samples, which could be considered as an improved maturation sign. Also, we could hypothesize that the bioprinting approach may have minimal impact on mRNA expression, primarily influencing the functional performance of the tissues. Therefore, further research is needed to fully understand how the bioprinting approach can affect muscle maturation at a cellular level.

## 3. Conclusions

The pluronic-assisted co-axial bioprinting (PACA-3D) hereby presented represents a novel and versatile approach to obtain individual bioengineered fibres with well-controlled homogeneity and width. Such individual filaments are of great interest for tissue engineering purposes, generating a functional fascicle-like structure that resembles the complexity of the native skeletal muscle tissue architecture. We explored two main hydrogel confinement methods: the first one based on chemical crosslinking of alginate upon extrusion thanks to the presence of CaCl2 within pluronic, and the second one based on physical repulsion between jellified gelatin and pluronic. Both strategies can be used in combination with any kind of biopolymer, such as fibrinogen, Matrigel®, GelMA, and others, making PACA-3D a universal 3D bioprinting method.

We demonstrated that our technology allows the fabrication of filaments of approximately 200 µm, which correspond to the oxygen diffusion limit, that do not fuse during the extrusion process. These filaments showed important advantages in the fabrication of skeletal muscle-based bioactuators when compared to the conventional extrusion-based 3D bioprinting. Probably due to the enhanced oxygen and nutrient diffusion, the myofibers obtained by PACA-3D clearly showed advanced maturation, resulting in 3 times stronger force output when compared to conventional bioactuators. Therefore, this bioprinting technique offers a valuable new strategy especially useful in skeletal muscle 3D bioengineering, paving the way towards an improved biomimetic model for basic developmental research or biomedical applications.

## 4. Materials and methods

### 4.1 Fabrication of the co-axial nozzles

The co-axial nozzles were manually fabricated using different types of commercially available nozzles and tips. The inner nozzle, where the cell-laden hydrogel passed through, was a 200-μm (G27) plastic conical nozzle (Optimum® SmoothFlow™ tapered tips, Nordson®, ref. 7018417). The luer lock of this nozzle was left free to be connected to the first bioprinting barrel, where the cell-laden hydrogel would be loaded. The outer nozzle that covered the inner one was a filtered P1000 pipette tip (Labclinics, ref. LAB1000ULFNL), cut approximately 5 cm from its end. The tip was trimmed to increase its diameter at the final point. Both nozzles were assembled and glued together. When the glue was dry and the assembly stable, a hot puncher was used to create a hole in the outer nozzle, at approximately 1.5 cm from its ending, with care to not create another whole in the internal nozzle. The secondary nozzle, where pluronic flowed, was inserted inside this hole. This secondary nozzle was a flexible polypropylene 800-μm nozzle (G18) from Nordson® (EFD® 7018138). A silicone tubing of 0.8 mm of diameter (Ibidi, ref. 10841) was attached to this external nozzle through a male elbow luer connector (Ibidi, ref. 10802). A 1.1-mm (19G) nozzle (B. Braun Sterican®, ref. 4657799) was inserted through the other end of the tubing, so that its luer lock connector could be connected to the second bioprinting barrel, containing pluronic acid.

### 4.2 Hydrogel fabrication

For the fabrication of the different combinations of hydrogels, the following materials were used: gelatin from porcine skin, type A (Sigma-Aldrich, G2500), fibrinogen from bovine plasma (Sigma-Aldrich, F8630) with thrombin from bovine plasma (Sigma-Aldrich, T4648) as crosslinker, Matrigel® Basement™ membrane matrix (Corning®, 354234), sodium alginate (Sigma-Aldrich, W201502), GelMA with lithium phenyl-2,4,6-trimethylbenzoylphosphinate (LAP) at 0.25% (wt/v) as photoinitiator (CELLINK®, LIK-3050V-1), collagen type I high concentration (Corning®, 354249), and Pluronic® F-127 powder (Sigma-Aldrich, P2443).

Pluronic was dissolved at concentrations ranging from 30-40% (wt/v) in Mili-Q water with CaCl2 (at molar concentrations ranging from 50 mM to 300 mM) under stirring in a refrigerator (4 °C) until fully dissolved. For hydrogels containing gelatin, alginate and/or GelMA in the same composition, these components were mixed together in PBS at the desired concentrations. If the hydrogel contained fibrinogen, this component was dissolved in PBS at a 2x concentration and then added to the hydrogel containing alginate and/or gelatin, also at a 2x concentration, in order to avoid pipetting this very viscous mixture. If the hydrogel also contained Matrigel®, the concentrations were also adjusted to achieve the desired concentrations reducing the need for pipetting (for instance, at 1:1:1 ratio, or 2:1:1 ratio, etc.).

### 4.3 Pluronic-Assisted Co-axial 3D Bioprinting

C2C12 myoblasts were harvested by a 0.25% (wt/v) Trypsin-0.53 mM EDTA solution, centrifuged at 300g for 5 min and the pellet re-suspended at a concentration of 5×106 cells/mL in the hydrogel mixture, at 37 °C. The cell-laden hydrogel was loaded into a 3-mL plastic syringe (Nordson®, ref. 7012085) coupled to the inner nozzle of the co-axial nozzle. Pluronic was loaded while cold into a secondary barrel and left at RT to gel beforehand. CELLINK® Inkredible+ 3D bioprinter was used to bioprint the hydrogel fibers. The cell-laden hydrogel was inserted in the first cartridge and the pluronic barrel to the second barrel, and all the nozzles connected as previously explained. The pressure for the pluronic barrel was kept between 250-350 kPa and adjusted manually, although these values are highly dependent on the diameter and length of the silicone tubing and the concentration of pluronic. For the hydrogel, the pressures were kept between 40-80 kPa, also adjusted manually, depending on the type of hydrogel. The designs were directly written in GCode with the help of the open-source software Slic3r (v. 1.2.9) and the bioprinter was controlled with RepetierHost (v. 2.0.5). After 3D bioprinting, if the hydrogel contained fibrinogen, a solution of 5 U/mL of thrombin was added to the Petri dish for 10 min in a refrigerator at 4 °C. At this point, pluronic would dissolve and fibrinogen would crosslink to form fibrin. After this, several washes with cold PBS were done to completely remove pluronic, GM supplemented with ACA was added and the constructs left in a cell incubator at 37 °C and 5% CO2 atmosphere. If the hydrogel contained Matrigel®, the as-printed constructs were left in an incubator for at least 30 min, without removing pluronic. After this, a cold solution of PBS (or thrombin, if fibrinogen was also in the mixture) was added to dissolve pluronic after several washes. If the chemical confinement method based on alginate was used, and this material wanted to be removed, a solution of 20 mM EDTA (Sigma-Aldrich, E6758) adjusted with NaOH to pH 7, was added to the Petri dish after crosslinking for 5 min.

### 4.4 Rheological characterization of pluronic acid

Rheological characterization of pluronic was performed using a Discovery HR-2 controlled-stress rheometer (TA Instruments) equipped with a Peltier steel cone geometry of 40 mm of diameter, 26 μm of truncation, and an angle of 1.019°. The Peltier element was set to 4 °C and 25 °C to demonstrate the behavior of the block co-polymer at low and room temperatures. In all experiments, the sample was left to acquire the desire temperature for 1 min. A flow ramp with shear rate from 100 1/s to 0.01 1/s was performed, in logarithmic mode with 600 s of duration per point, with a pre-conditioning to the temperature of 30 s and pre-shear of 3 rad/s for 10 s.

### 4.5 Cell culture, differentiation, and electrical stimulation

C2C12 mouse myoblasts were purchased from ATCC and maintained in growth medium (GM) consisting of high glucose Dullbecco’s Modified Eagle’s Medium (DMEM; Gibco®) supplemented with 10% Fetal Bovine Serum (FBS), 200 nM L-Glutamine and 1% Penicillin/Streptomycin, in a 37 °C and 5% CO2 atmosphere. Cells were passaged before reaching 80% confluence in Corning® T-75 flasks. For cell differentiation and myotube formation after the bioprinting process, GM was substituted by DM, consisting of high glucose DMEM containing 10% v/v Horse Serum (Gibco®), 200 nM L-Glutamine (Gibco®), 1% v/v Penicillin-Streptomycin (Gibco®), 50 ng/ml IGF-1 (Sigma-Aldrich) and 1 mg/ml 6-aminocaproic acid (ACA, Sigma-Aldrich). 3D-bioprinted fibers were stimulated with a set of carbon-made electrodes attached to the cover of a Petriunder an inverted microscope (Leica’s DMi8) with pulses of 2 ms and 1 V/mm. Analysis of the contractions was performed with a home-made Python algorithm based on computer vision techniques that computed the distance between frames of a selected ROI by applying an L2 norm to the image pixels.

### 4.6 Cell viability and MyHCII immunofluorescence

Cell viability was analyzed by the dual-fluorescence Live/Dead® Viability/Cytotoxicity kit for mammalian cells (Life Technologies), following the manufacturer’s instructions. Fluorescently labelled fibers were imaged under a Leica DMi8 inverted fluorescence microscope equipped with a 37 °C and 5% CO2 chamber, using a 20× air objective. The percentages of live and dead cells were calculated by using ImageJ software ver.1.47q (National Institutes of Health, Bethesda, MD). For immunostaining, 3D-bioprinted constructs were washed twice in PBS and fixed by incubating them with a 3.7% v/v paraformaldehyde in PBS solution for 15 min at RT, followed by three washes in PBS. Then, cells were permeabilized by using 0.2% v/v Triton-X-100 in PBS. After two washes in PBS, the constructs were incubated with 5% w/v Bovine Serum Albumin (BSA) in PBS (PBS-BSA) to block unspecific bindings. Then, bioprinted structures were incubated for 2 h at RT and dark conditions with a 1/400 dilution of Alexa Fluor®488-conjugated Anti-Myosin Heavy Chain II antibody (eBioscience, 53-6503-82) in 5% w/v PBS-BSA. The unbound antibodies were washed out with PBS, and cell nuclei were counterstained with 1 μl/mL Hoechst 33342 (Life technologies). Finally, samples were washed twice in PBS, and they were stored at 4 °C until their analysis. Fluorescently immunostained fiber constructs were imaged under a Zeiss LSM 800 confocal scanning laser microscope (CSLM), with a diode laser at 488 nm and 405 nm excitation wavelength for Myosin Heavy Chain II and cell nuclei.

The orientation and alignment of nucleus and myotubes was analyzed via Image J with the plugin Directionality. The histograms and angle dispersion values were obtained from the Fast Fourier Transform (FFT) of the immunostaining images.

### 4.7 Force Measurement

Evaluation of the force output of muscle tissues was performed as reported by Mestre et al.[10] Briefly, the two-posts system where the tissue is directly printed and maintained during differentiation was used as a force measurement platform. Muscle contraction was induced with pulse electrical stimulation by applying 15V at 1 Hz frequency of 2 ms. Contractile tissues generated a force against the flexible posts which bend synchronized with the applied electrical pulse. The recording of the set up was performed with a Thunder Leica microscope where the bending of the post can be easily tracked. Then, to estimate the force exerted against the post, Eurler-Bernoulli’s beam bending equation was used, that is, using the following equation:

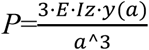

Where *P* is the applied force, *E* is the Young’s modulus, *Iz* is the second moment of area of the post around the z axis, a is the height from which the tissue is exerting the force and *y(a)* is the displacement of the post at such height.

### 2.8 RNA extraction and Real Time Quantitative PCR (RT-qPCR)

To evaluate the mRNA expression of myogenic markers such as MyoD, myogenin and MyHCI, the total RNA content from 3 biological replicates of muscle tissues printed with the coaxial and conventional approach were extracted at days 0, 4, 9 and 12 of differentiation. RNeasy mini-Kit (Qiagen, 74134) was used for this purpose. RNA was quantified by absorbance at 260 nm in a nano-drop spectrophotometer (ND-1000, Nano-Drop) and converted into cDNA using the ReverAid First Strand cDNA Synthesis Kit (Thermo Scientific™, K1622). Finally, 500 ng of cDNA was be mixed with the corresponding forward and reverse primers together with the PowerUp SYBR Green Master Mix (Applied Biosystems, A25742), following manufacturer’s instructions, and RT-qPCR were carried out in an Applied Biosystems StepOnePlus Real-Time PCR (Applied Biosystems, 4376600). The list of primers can be found in supplementary info. All genes were normalized to the expression levels of GAPDH.

## Data availability statement

The data that support the findings of this study are available from the corresponding author upon request.

## Acknowledgements

R.M. would like to thank UK Research and Innovation (UKRI) funding (grant refs ES/X006255/1 and MR/S032711/1). M. G. acknowledges the financial support from the Spanish Ministry of Science through the Ramon y Cajal Grant No. RYC2020-945030119-I. and Unidades de Excelencia María de Maeztu» 2021 CEX2021-001202-M. J.F. and SS acknowledges the financial support from the European Union’s Horizon Europe research and innovation programme under grant agreement No. 101070328 (Biomeld). S.S. acknowledges CERCA program by the Generalitat de Catalunya, the Secretaria d’Universitats i Recerca del Departament d’Empresa i Coneixement de la Generalitat de Catalunya through the project 2021 SGR 01606, and the "Centro de Excelencia Severo Ochoa", funded by Agencia Estatal de Investigación (CEX2018-000789-S).N.R. is thankful for the financial support from the European Research Council (ERC) under the European Union’s Horizon 2020 research and innovation programme (grant agreement No 866348, i-NanoSwarms) and the Spanish Ministry of Science (grants PID2021-128417OB-I00 and RETI2018-098164-B-I00 funded by MCIN/AEI/10.13039/501100011033). T.P. thanks the Irène Curie Fellowship.

---

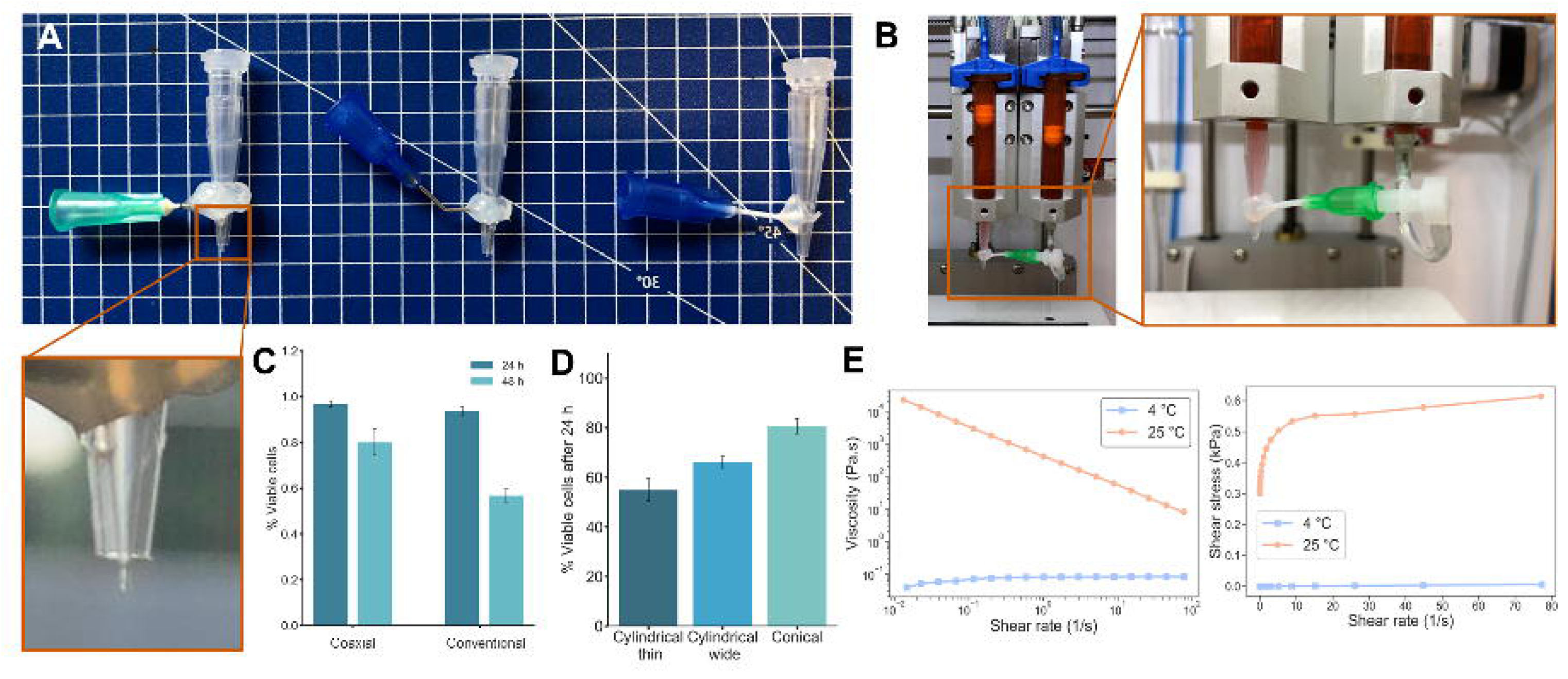

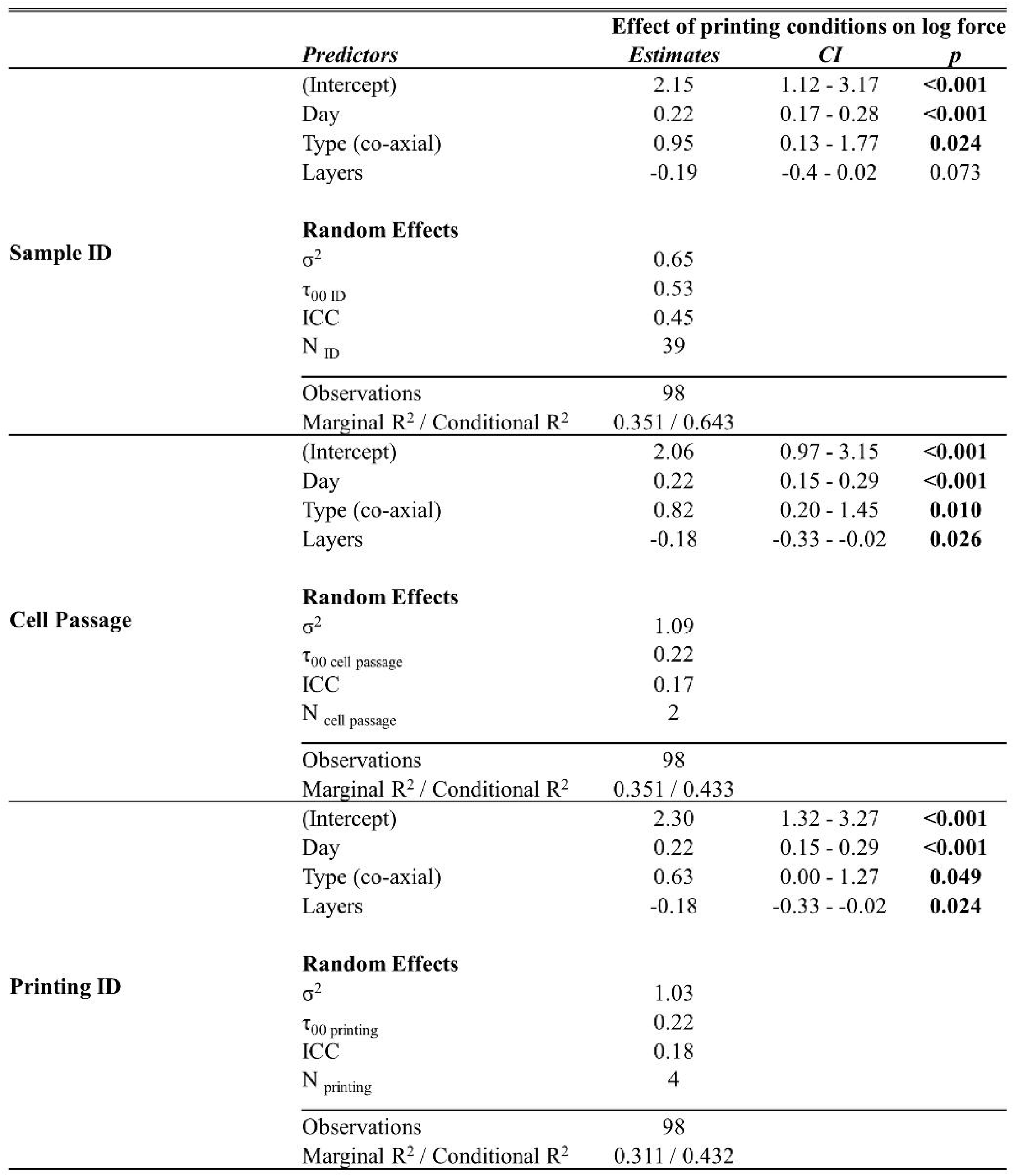

|  |  | Effect of printing conditions on log force |  |  |
| --- | --- | --- | --- | --- |
|  | Predictors | Estimates | CI | p |
| Sample ID | (Intercept) | 2.15 | 1.12 - 3.17 | <0.001 |
|  | Day | 0.22 | 0.17 - 0.28 | <0.001 |
|  | Type (co-axial) | 0.95 | 0.13 - 1.77 | 0.024 |
|  | Layers | -0.19 | -0.4 - 0.02 | 0.073 |
|  | Random Effects |  |  |  |
| | $\sigma^2$ | 0.65 | | |
| | $\tau_{00 \text{ ID}}$ | 0.53 | | |
|  | ICC | 0.45 |  |  |
| | $N_{\text{ID}}$ | 39 | | |
|  | Observations | 98 |  |  |
| Marginal R<sup>2</sup> / Conditional R<sup>2</sup> |  | 0.351 / 0.643 |  |  |
| Cell Passage | (Intercept) | 2.06 | 0.97 - 3.15 | <0.001 |
|  | Day | 0.22 | 0.15 - 0.29 | <0.001 |
|  | Type (co-axial) | 0.82 | 0.20 - 1.45 | 0.010 |
|  | Layers | -0.18 | -0.33 - -0.02 | 0.026 |
|  | Random Effects |  |  |  |
| | $\sigma^2$ | 1.09 | | |
| | $\tau_{00 \text{ cell passage}}$ | 0.22 | | |
|  | ICC | 0.17 |  |  |
| | $N_{\text{cell passage}}$ | 2 | | |
|  | Observations | 98 |  |  |
| Marginal R<sup>2</sup> / Conditional R<sup>2</sup> |  | 0.351 / 0.433 |  |  |
| Printing ID | (Intercept) | 2.30 | 1.32 - 3.27 | <0.001 |
|  | Day | 0.22 | 0.15 - 0.29 | <0.001 |
|  | Type (co-axial) | 0.63 | 0.00 - 1.27 | 0.049 |
|  | Layers | -0.18 | -0.33 - -0.02 | 0.024 |
|  | Random Effects |  |  |  |
| | $\sigma^2$ | 1.03 | | |
| | $\tau_{00 \text{ printing}}$ | 0.22 | | |
|  | ICC | 0.18 |  |  |
| | $N_{\text{printing}}$ | 4 | | |
|  | Observations | 98 |  |  |
| Marginal R<sup>2</sup> / Conditional R<sup>2</sup> |  | 0.311 / 0.432 |  |  |

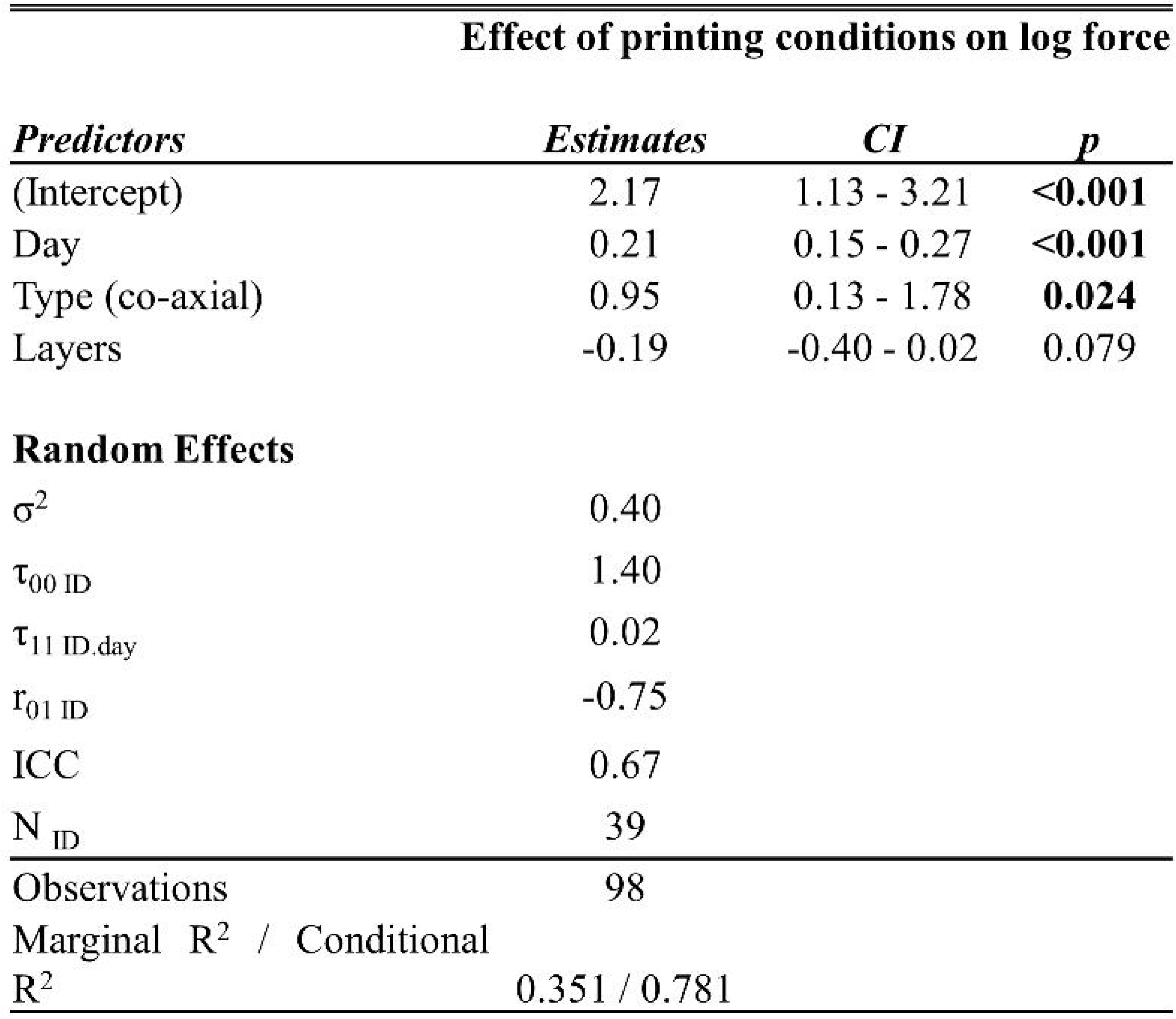

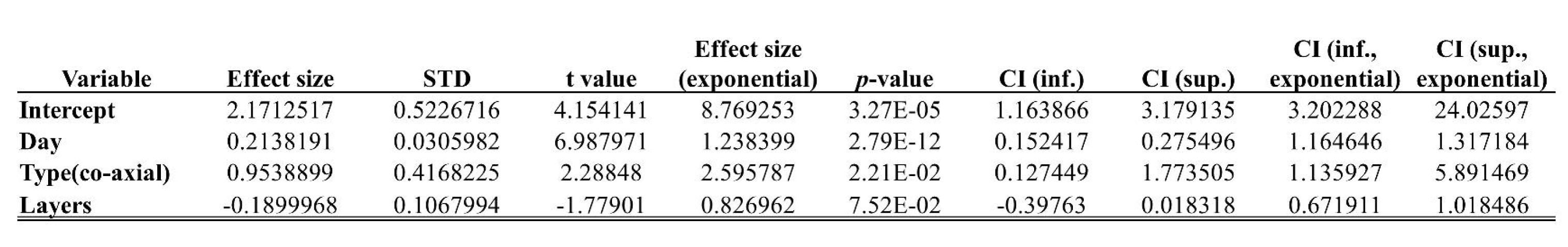

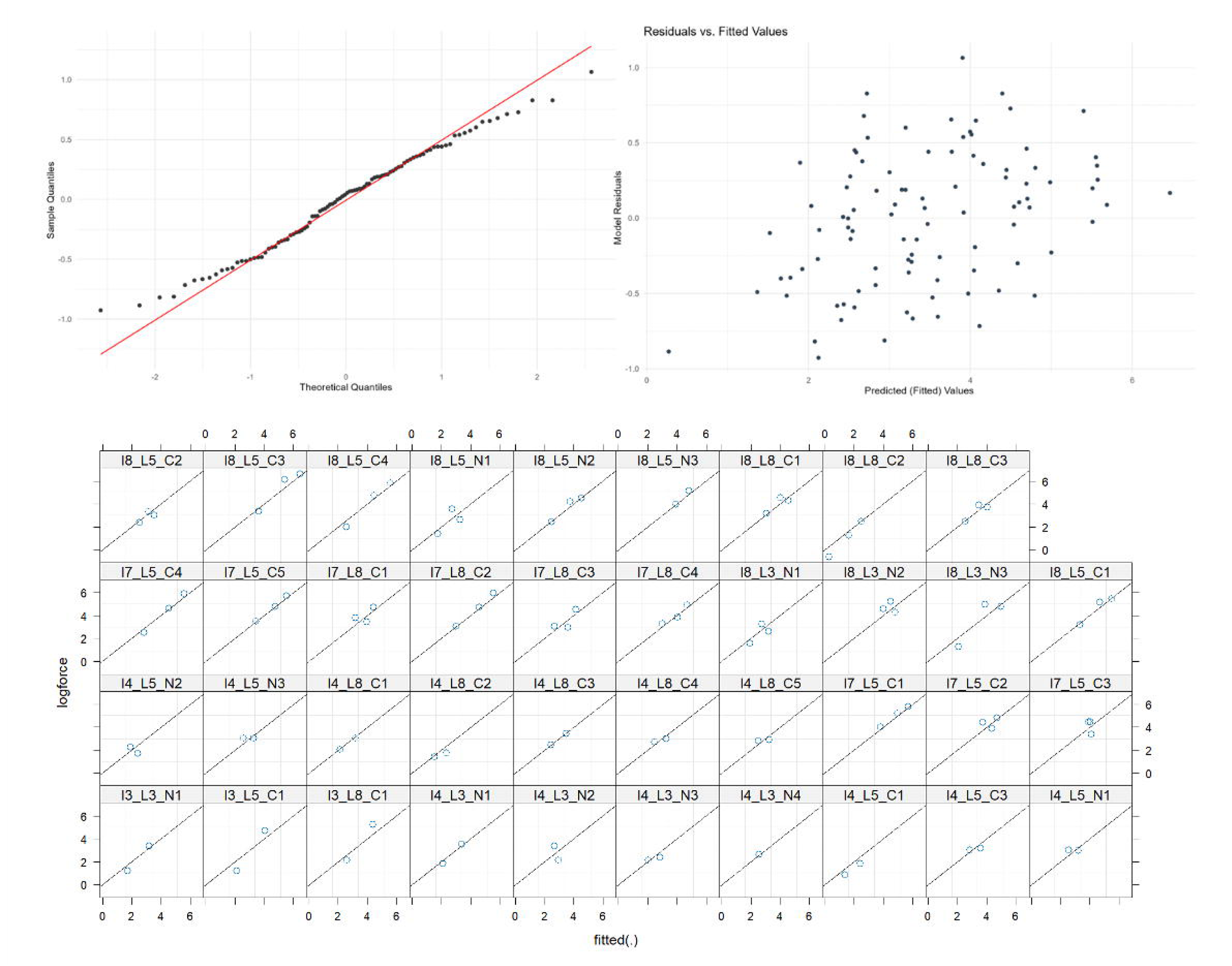

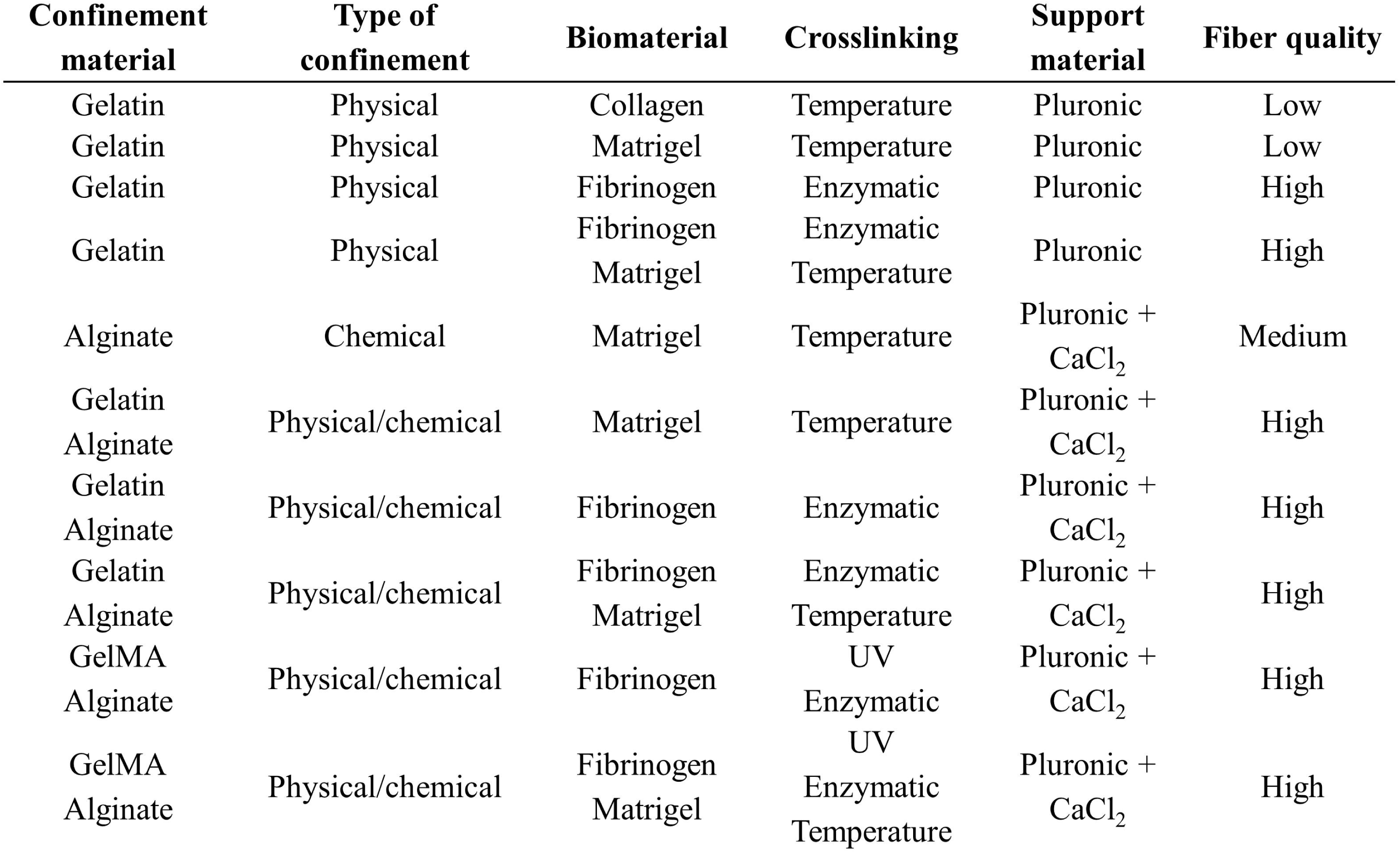

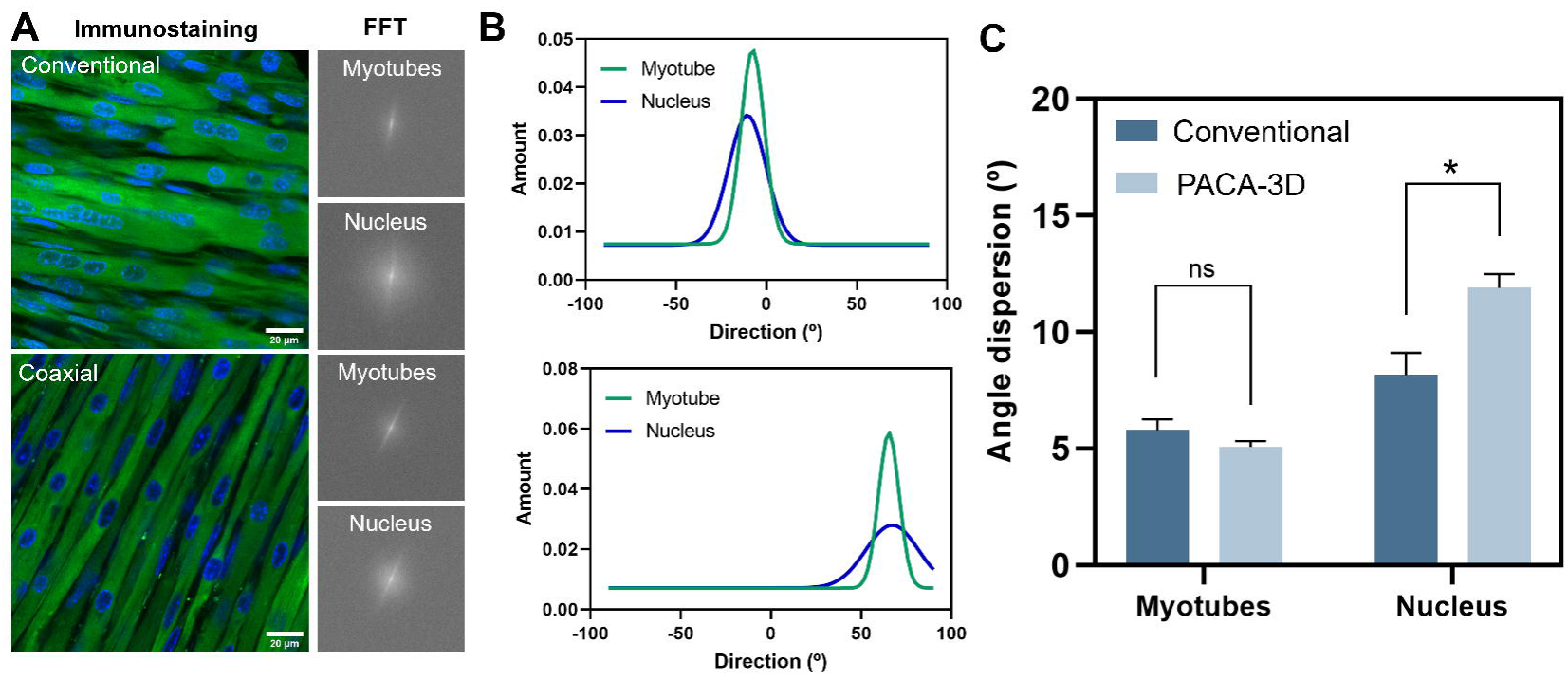

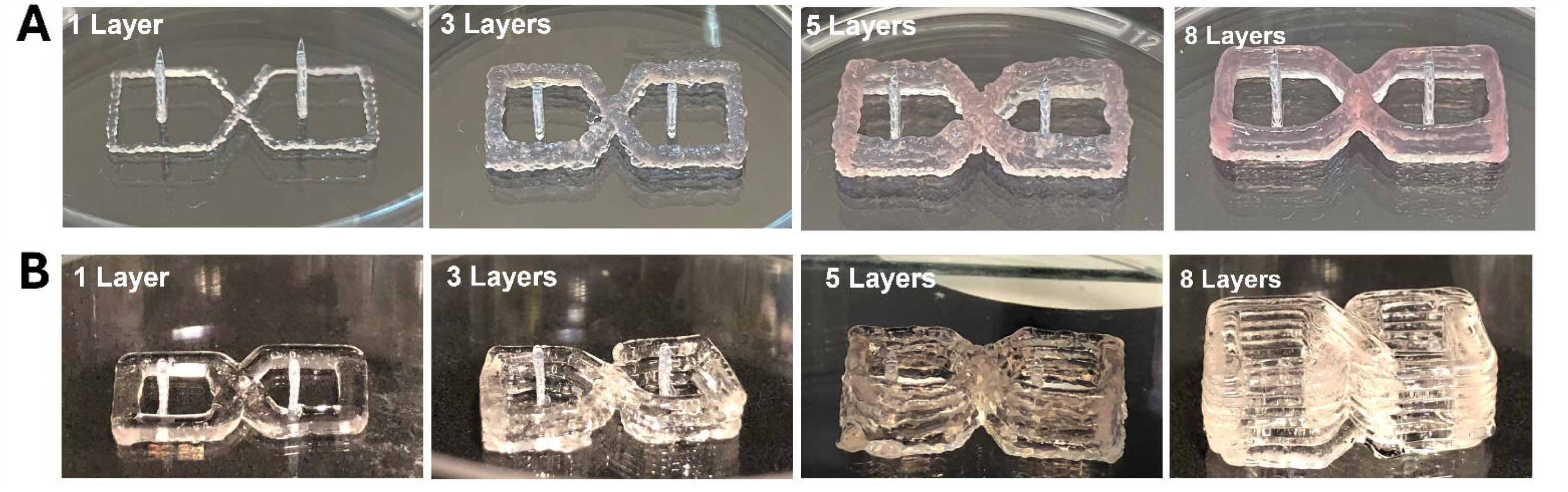

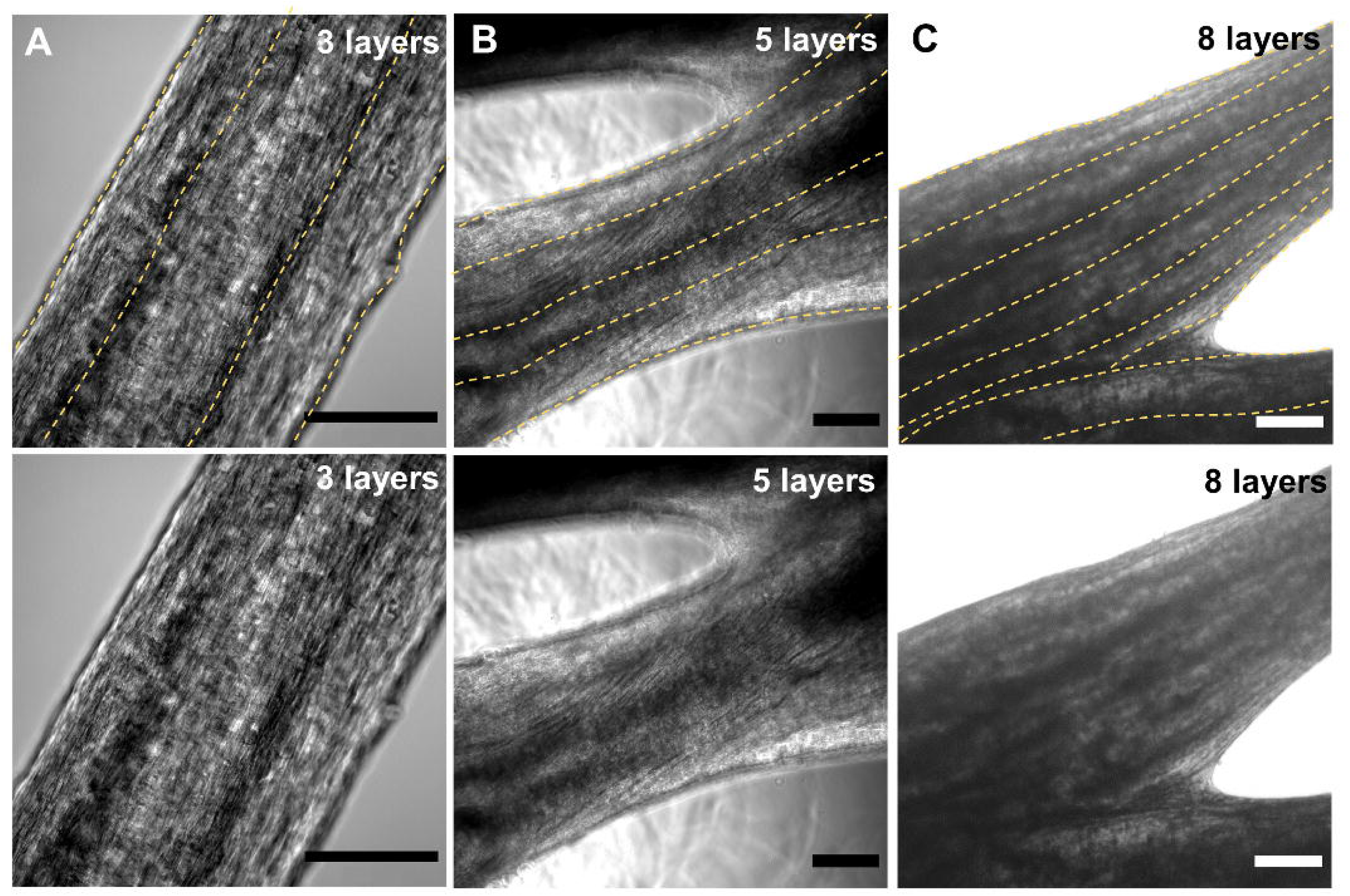

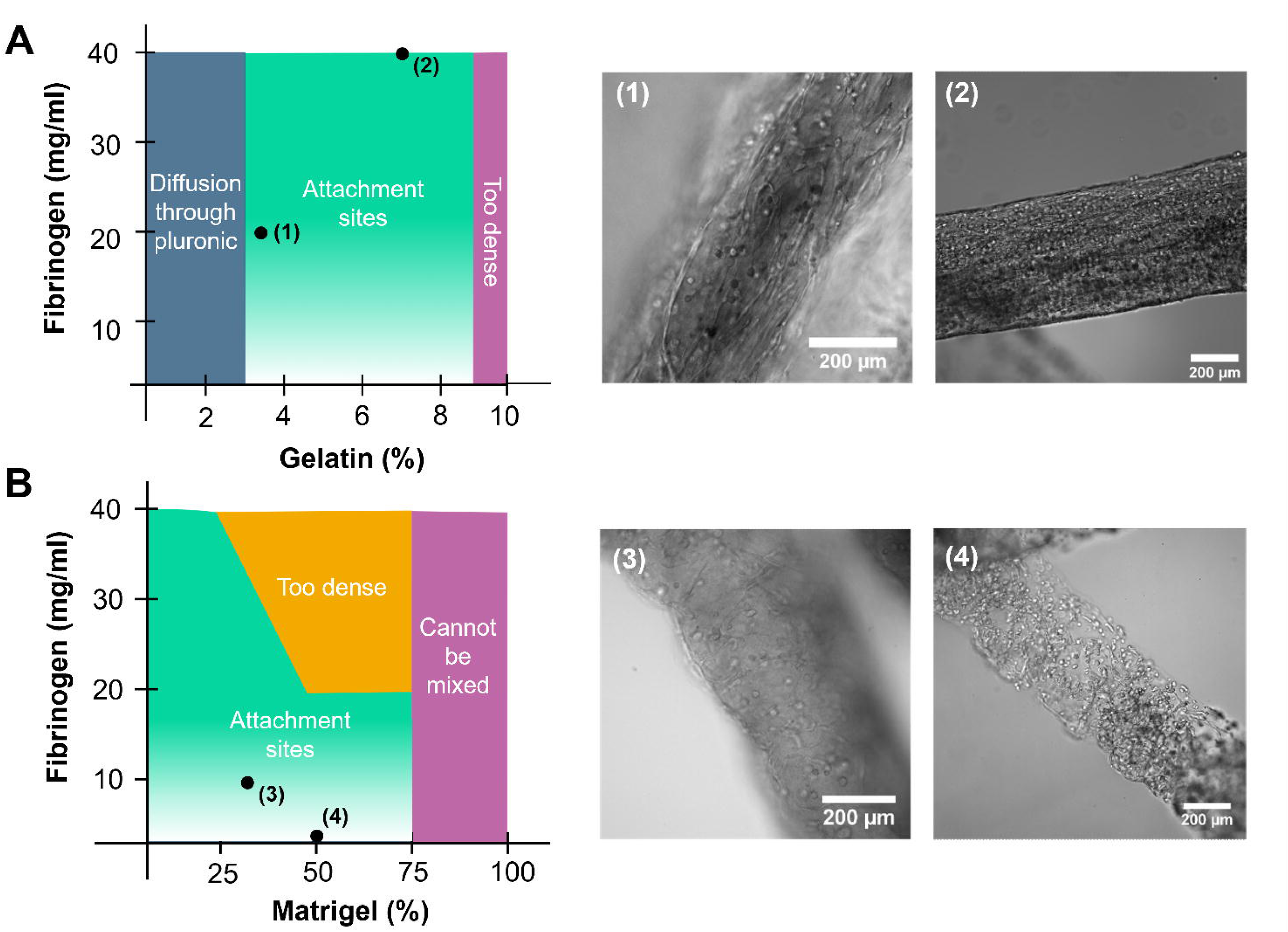

