## Supplementary Information text for "Bioengineering Fascicle-like Skeletal Muscle Bioactuators via Pluronic-Assisted Co-axial 3D Bioprinting"

#### 1.Comparison of different biomaterials, confinements strategies and crosslinking process.

The column fiber quality is influenced by both the homogeneity of the printing and the stability of the fibers. For instance, physical confinement of a hydrogel only containing Matrigel, or collagen might give homogeneous fibers if the concentration of gelatin is high enough, but their stability is low, as their stiffness of the construct after temperature crosslinking is low. Therefore, thin and homogeneous fibers can be printed with this combination, but they are easily broken. Adding fibrinogen to the mixture increases their long-term stability, as fibrin is more robust, creating high quality fibers. A similar effect occurs with the chemical confinement method. If the main biomaterial is too liquid, like Matrigel, it will diffuse through the pluronic before the Ca-crosslinking of alginate can form homogeneous fibers, unless the concentration of alginate is increased to accelerate this process. Since alginate does not have cell attachment motives, it is advisable to keep its concentration as low as possible; in that case, however, the quality of the fiber will not be high. One of the best strategies to follow is the combination of both methods by adding a small amount of gelatin or GelMA to alginate, using a hybrid chemical-physical method of confinement. With this combination, as can be seen in the table, the quality of the fibers is much improved and virtually any material can be used with it.

Collagen and Matrigel, which are crosslinked slowly at 37 °C for approximately 30 min in an irreversible manner, deserve special consideration. Collagen is one of the main components of many tissue's ECM and, in particular, of skeletal muscle tissue, making it especially interesting for

3D bioengineering applications [58]. Also, Matrigel is one of the most widely used basement membrane matrices for 2D and 3D cultures, since it is rich in collagen and many other ECM proteins, but shares the same difficulties. However, their irreversible and low temperature-dependent crosslinking makes difficult their bioprinting, as opposed to gelatin, which has reversible crosslinking. Both materials are liquid at room temperature and cannot be 3D-bioprinted by pneumatic extrusion, but if they are crosslinked at 37 °C, they are also too stiff to be extruded. Although the most promising approaches in the literature have dealt with the mixture of alginate and collagen to achieve proper extrusion [59], better strategies are necessary in the field. One of the main difficulties with these approaches comes from the slow crosslinking of these materials. Because of this, the printed construct can easily lose its shape before the crosslinking has taken place. For that reason, collagen or Matrigel have been mainly used with casting molds, which can retain the shape of the construct until the hydrogel is fully crosslinked [47]. With the pluronic-assisted co-axial printing method, however, these materials can be protected during crosslinking at 37 °C, avoiding its diffusion in the media. As pluronic does not dissolve at this temperature, the fibers can be incubated in physiological temperatures for 30-45 min until Matrigel or collagen have been crosslinked, and then pluronic can be removed with cold PBS. Both confinement methods (either chemical or physical) could potentially be used in combination with this type of hydrogel, followed a tertiary crosslinking with fibrinogen to improve the mechanical stability of the fibers.

**Table S1:** Relation of different strategies to obtain thin, independent fibers, combining different biomaterials, confinement strategies and crosslinking methods.

| Confinement material | Type of confinement | Biomaterial | Crosslinking | Support material | Fiber quality |
|---|---|---|---|---|---|
| Gelatin | Physical | Collagen | Temperature | Pluronic | Low |
| Gelatin | Physical | Matrigel | Temperature | Pluronic | Low |
| Gelatin | Physical | Fibrinogen | Enzymatic | Pluronic | High |
| Gelatin | Physical | Fibrinogen<br>Matrigel | Enzymatic<br>Temperature | Pluronic | High |
| Alginate | Chemical | Matrigel | Temperature | Pluronic + $CaCl_2$ | Medium |
| Gelatin<br>Alginate | Physical/chemical | Matrigel | Temperature | Pluronic + $CaCl_2$ | High |
| Gelatin<br>Alginate | Physical/chemical | Fibrinogen | Enzymatic | Pluronic + $CaCl_2$ | High |
| Gelatin<br>Alginate | Physical/chemical | Fibrinogen<br>Matrigel | Enzymatic<br>Temperature | Pluronic + $CaCl_2$ | High |
| GelMA<br>Alginate | Physical/chemical | Fibrinogen | UV<br>Enzymatic | Pluronic + $CaCl_2$ | High |
| GelMA<br>Alginate | Physical/chemical | Fibrinogen<br>Matrigel | UV<br>Enzymatic<br>Temperature | Pluronic + $CaCl_2$ | High |

### 2. Optimization of the co-axial needles

Different types of co-axial nozzles were manually fabricated, as shown in figure S1. These needles consisted of a primary nozzle, where the cell-laden hydrogel would pass through, and a secondary nozzle through which pluronic would be extruded. The primary nozzle was always a 200-µm conical plastic nozzle to ensure viability of the printed cells. The challenge, therefore, was to assemble the secondary external needle. For this purpose, a disposable micropipette P1000 tip was cut off and the conical nozzle introduced inside of it, gluing them together. A heated puncher was used to make a hole at one of the sides of the micropipette tip, leaving room for the insertion of the secondary nozzle that would allow the flow of pluronic around the primary conical nozzle.

The zoomed-in picture in Figure S1A displays the state of the tip after assembly, where we can see how the inner nozzle protrudes surrounded by the outer one.

Several secondary nozzles for pluronic were considered: a straight steel nozzle, a bent nozzle and a flexible cylindrical nozzle (figure S1A). A straight steel nozzle (left) was not compatible with the bioprinting setup, as it would crash with the walls of the Petri dish during printing. A bent steel nozzle would partially solve this issue (center), but it would leave less room for versatility, and it might still collide with the walls of the dish. Finally, a flexible nozzle (right) solved the problem, as it would adapt to the geometry around the Petri dish avoiding collisions during the printing process. The assembled co-axial nozzle was adapted to 3D bioprinting setup to control both flows of materials (figure S1 B). The inner nozzle was directly connected to the first cartridge and would apply pressure control from the first printhead. The external nozzle was connected through a silicone tubing to the second printhead to control the pluronic flow.

Straight cylindrical co-axial needles are easier to assemble due to their simpler shape, but they pose more damage for cells due to increased shear stress. Figure S1(D**)** shows a Live/Dead characterization that proves this statement. We 3D-bioprinted a cell-laden hydrogel with the pluronic-based co-axial system, considering three different primary nozzles: thin (700 um diameter), wide (840 um diameter approximately) and a plastic conical nozzle of 200 μm of diameter at the tip. After 24 h, cell viability was higher for the cells printed using the conical nozzle, despite having a smaller nozzle diameter.

Finally, rheological characterization of pluronic acid was performed at low and room temperatures to investigate the use of this material as a sheath (figure S1E). At 4 °C, pluronic behaves as a Newtonian liquid, with no yield stress and a constant viscosity over the range of shear rate tested. At room temperature, however, it shows a strong shear-thinning effect with a finite yield stress and a decreasing viscosity for higher shear rates, explaining its smooth printability and shape retaining.

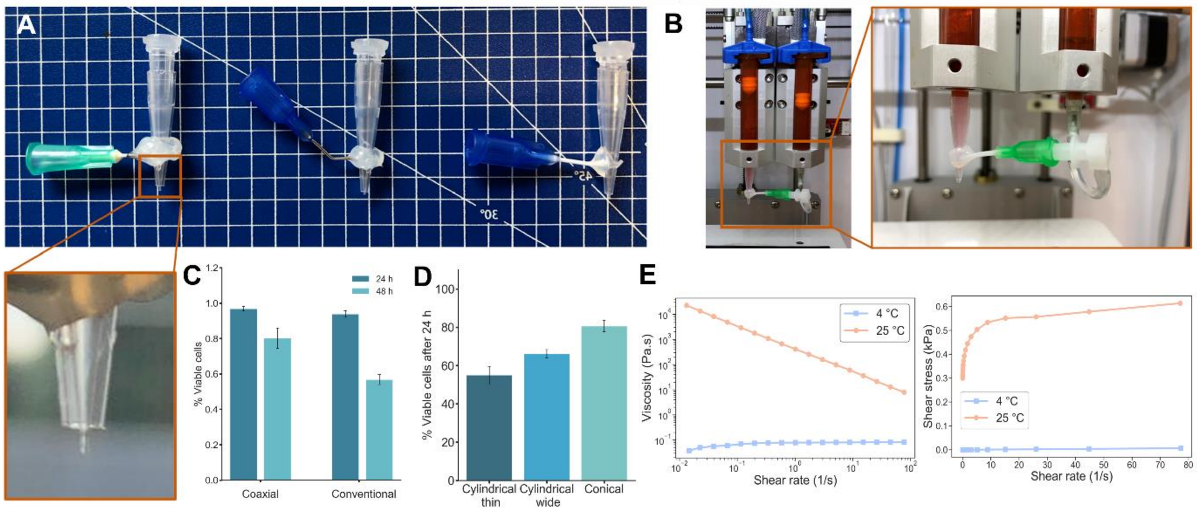


**Figure S1. Optimization of the PACA-3D system.** (A) Several types of co-axial nozzles based on conical needles were considered, offering different degrees of versatility by modifying the outer needle to obtain different angles of insertion or flexibility. The tip of a co-axial needle can be observed in the zoomed-in image (orange square), showing how the outer nozzle surrounds the inner one.  (B) Both cartridges of the 3D bioprinter are used to control the flow of both materials, connecting the outer needle to the second cartridge with a microfluiding tubing. (C) Cell viability after co-axial and conventional printing. When the tissue is fabricated with conventional printing, the width of the hydrogel cannot be properly controlled, yielding thicker fibers. Besides having similar viability after 24 h, there is a higher decrease of variability after 48 h. (D) Cell viability after 24 h shows that conical needles provide less cell damage due to less amount of shear stress. (E) Rheological characterization of Pluronic acid at 35% (wt/v) for two different temperatures. At low temperature, Pluronic behaves as a Newtonian liquid with constant viscosity. At room temperature, it shows a strong shear-thinning behavior.

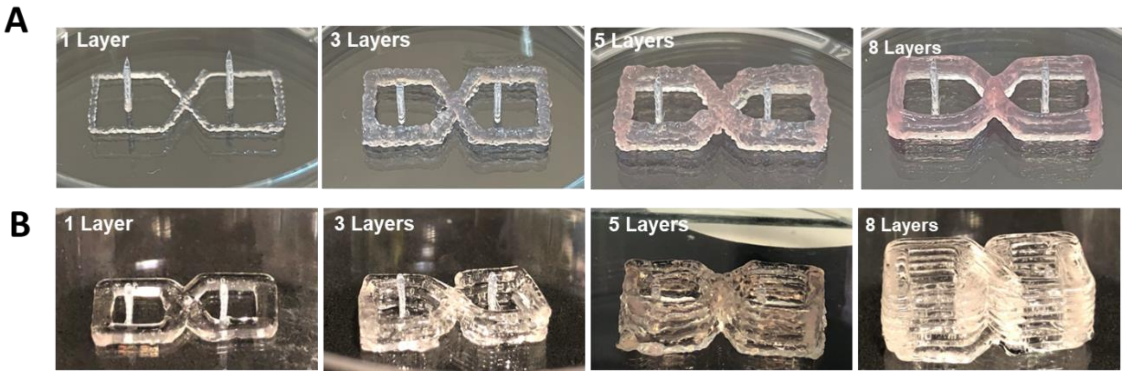


**Figure S2**. Images of 3D printed constructs using conventional printing (A) and PACA-3D printing (B).

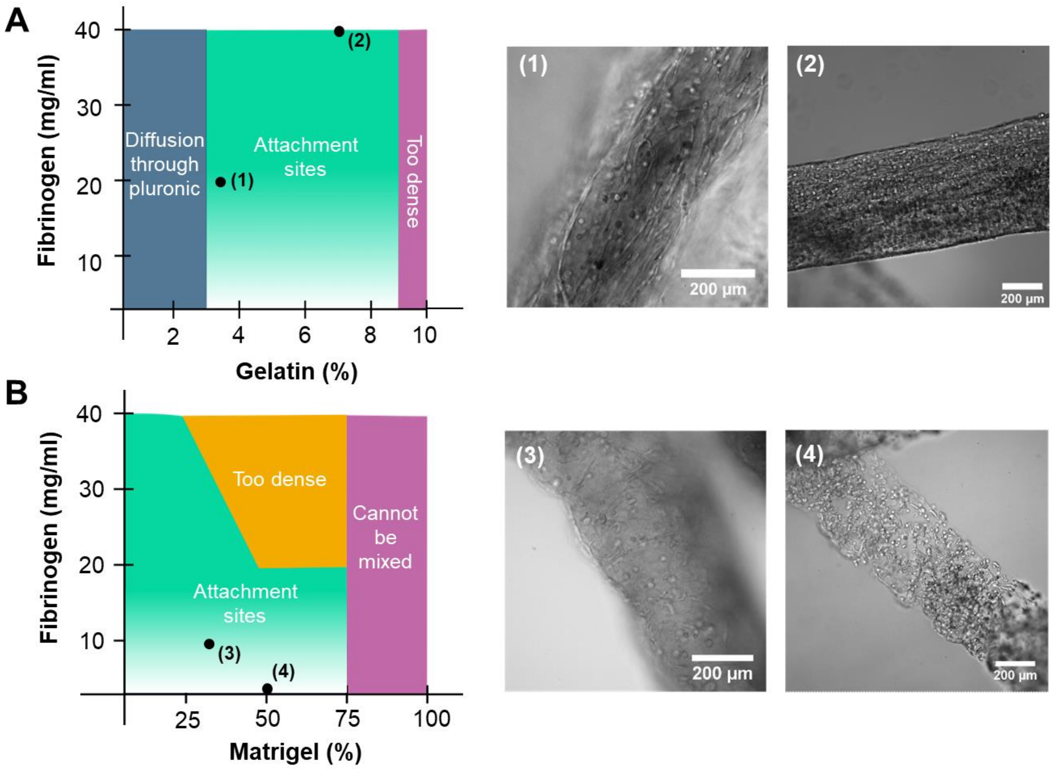


**Figure S3.** Ranges of application of materials according to their biocompatibility. Note that the different Matrigel-fibrinogen bioinks (lower plot) contain 7% (wt/v) of gelatin. Microscope images show attachment and alignment of the cells after 1 day of culture

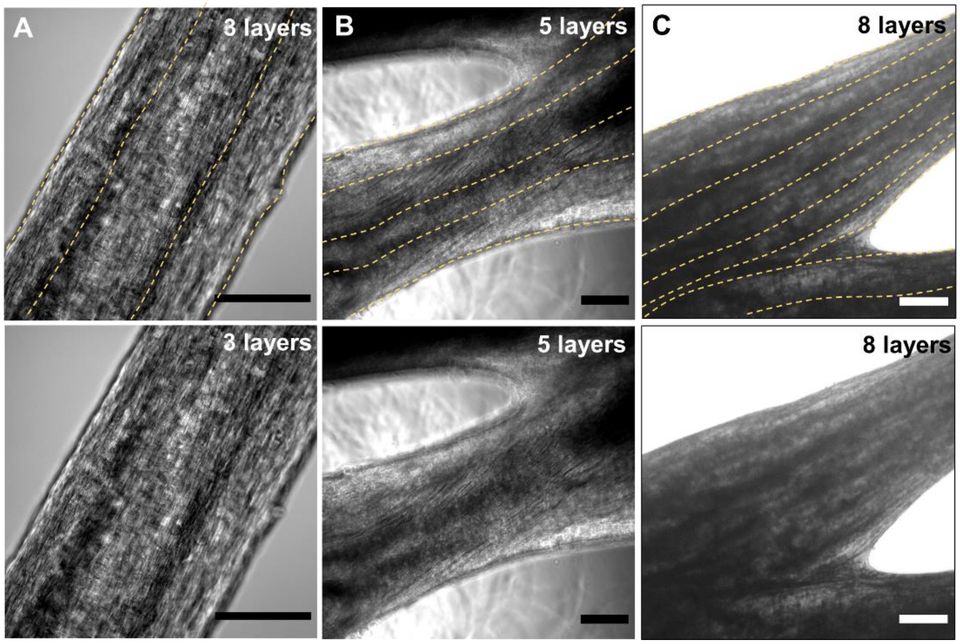


**Figure S4.** Representative examples of skeletal muscle tissue construct 3D bioprinted with the Pluronic-assisted co-axial method, with dashed lines representing separation of the tissue stripes, which were indicated by the changes in

bright field light intensity, for (A) 3 layers, (B) 5 layers and (C) 8 layers. Not all layers can be observed as some might be located below other layers. Scale bar: 200 µm.

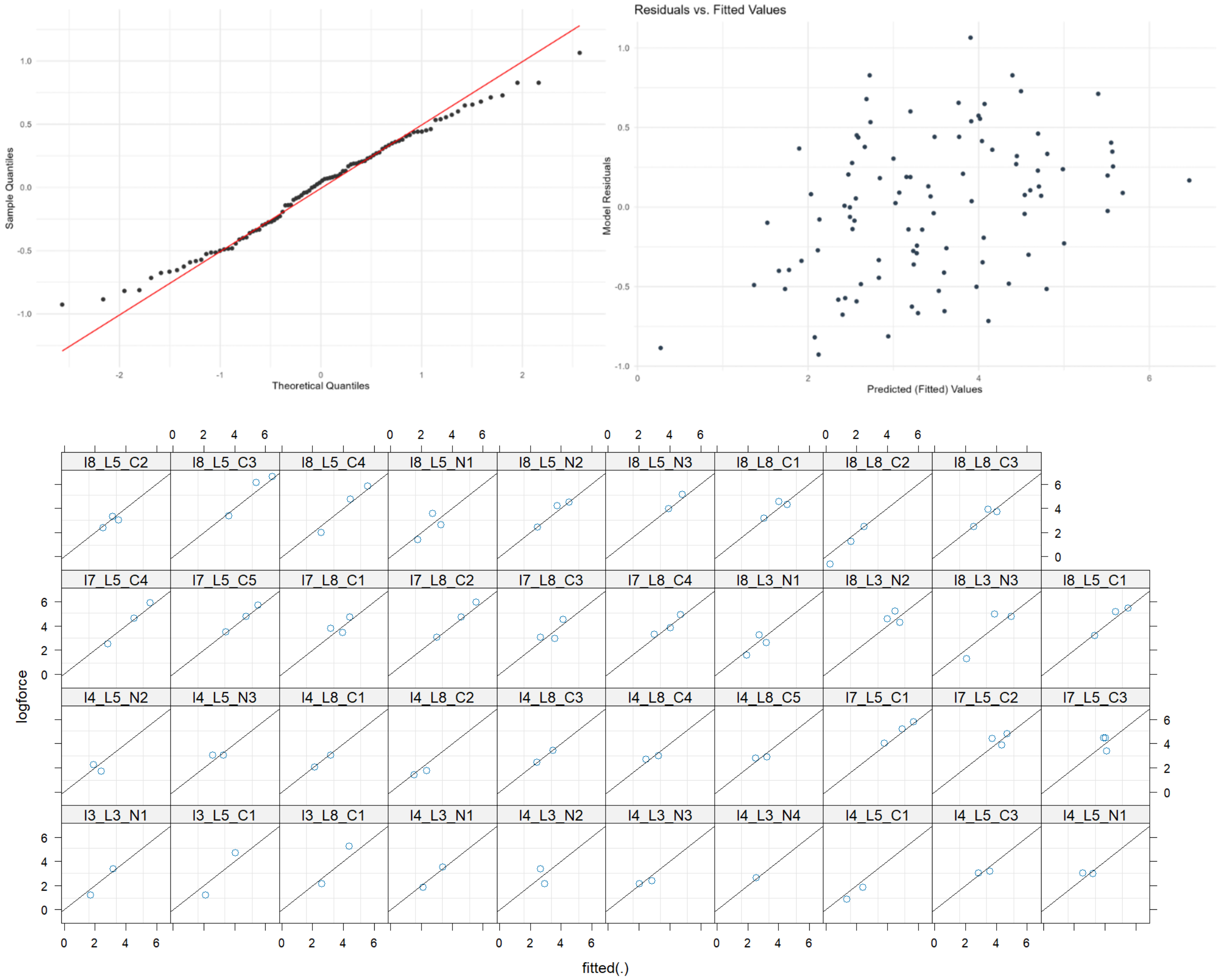


**Figure S5**. Diagnostic plots for the linear mixed-effects model. The top left displays a QQ-plot of residuals with the theoretical quantiles plotted against the sample quantiles. The residuals closely follow the line, especially at the center, indicating a close to normal distribution. The top right presents the overall residuals vs. fitted values, highlighting potential deviations from homoscedasticity. There are no visible patterns in this data, indicating that the data is homoscedastic. The plots at the bottom show residuals vs. fitted values by ID, allowing for a more detailed inspection of individual variance and potential patterns or outliers. None of the values seem to deviate strongly from the expected.

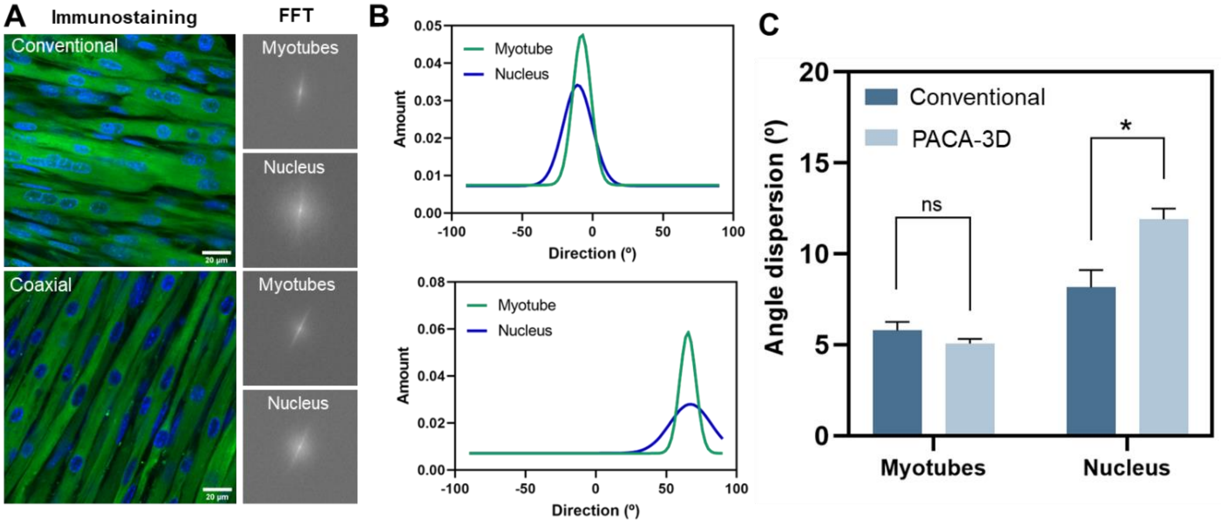


**Figure S6.** Analysis of the myotube and nucleus alignment on PACA-3D and conventional bioprinted muscles at day 13 of differentiation. (A) Left: images of the immunofluorescence staining of Myosin 4 (i.e Myosin Heavy Chain IIb) in green and cell nucleus (i.e. Hoechst) in blue; right: Fast Fourier transform (FFT) images obtained from the immunostaining images, using Image J. (B) Directionality histograms showing the level of alignment (i.e., orientation) of both myotubes and nucleus. The wider the normal distribution, the less oriented or aligned are muscle cells. Top and bottom graphs correspond to conventional and PACA-3D printed samples, respectively. C) Graph displaying the values of angle dispersion (º) of both myotubes and nucleus. Higher values of angle dispersion indicate less orientation and alignment of the muscle fibers and nucleus. Nucleus from conventionally printed tissues present significant lower values of angle dispersion in comparison with the nucleus from PACA-3D printed samples, probably because of the nucleus dispersion and disposition in the center of the myofiber, which suggests that they are in a more mature phase than PACA-3D samples. All values are represented as mean ± SEM. $*p < 0.05$. A Student T-test was performed for the analysis of nucleus and myotubes alignment (N=7-8 images from a sample printed with conventional and PACA-3D).

**Table S2**. Comparison of the random effects models in Sample ID, cell passage and Printing ID. Models calculated using the *lmer*() function from the *lme4* library (v. 1.1-34) in R (v. 4.3.1). 95% confidence intervals are calculated using the *wald* approximation.

| | | **Effect of printing conditions on log force** | | |
|---|---|---|---|---|
| | ***Predictors*** | ***Estimates*** | ***CI*** | ***p*** |
| | (Intercept) | 2.15 | 1.12 - 3.17 | **<0.001** |
| | Day | 0.22 | 0.17 - 0.28 | **<0.001** |
| | Type (co-axial) | 0.95 | 0.13 - 1.77 | **0.024** |
| | Layers | -0.19 | -0.4 - 0.02 | 0.073 |
| **Sample ID** | **Random Effects** | | | |
| | $\sigma^2$ | 0.65 | | |
| | $\tau_{00\ ID}$ | 0.53 | | |
| | ICC | 0.45 | | |
| | $N_{ID}$ | 39 | | |
| | Observations | 98 | | |

| | | | | |
|---|---|---|---|---|
| | Marginal $R^2$ / Conditional $R^2$ | 0.351 / 0.643 | | |
| Cell Passage | (Intercept) | 2.06 | 0.97 - 3.15 | **<0.001** |
| | Day | 0.22 | 0.15 - 0.29 | **<0.001** |
| | Type (co-axial) | 0.82 | 0.20 - 1.45 | **0.010** |
| | Layers | -0.18 | -0.33 - -0.02 | **0.026** |
| | **Random Effects** | | | |
| | $\sigma^2$ | 1.09 | | |
| | $\tau_{00\ cell\ passage}$ | 0.22 | | |
| | ICC | 0.17 | | |
| | $N_{cell\ passage}$ | 2 | | |
| | Observations | 98 | | |
| | Marginal $R^2$ / Conditional $R^2$ | 0.351 / 0.433 | | |
| Printing ID | (Intercept) | 2.30 | 1.32 - 3.27 | **<0.001** |
| | Day | 0.22 | 0.15 - 0.29 | **<0.001** |
| | Type (co-axial) | 0.63 | 0.00 - 1.27 | **0.049** |
| | Layers | -0.18 | -0.33 - -0.02 | **0.024** |
| | **Random Effects** | | | |
| | $\sigma^2$ | 1.03 | | |
| | $\tau_{00\ printing}$ | 0.22 | | |
| | ICC | 0.18 | | |
| | $N_{printing}$ | 4 | | |
| | Observations | 98 | | |
| | Marginal $R^2$ / Conditional $R^2$ | 0.311 / 0.432 | | |

**Table S3:** Random effects model allowing random effects in sample ID and differentiation day. Model calculated using the *lmer*() function from the *lme4* library (v. 1.1-34) in R (v. 4.3.1). 95% confidence intervals are calculated using the *wald* approximation.

| | **Effect of printing conditions on log force** | | |
|---|---|---|---|
| ***Predictors*** | ***Estimates*** | ***CI*** | ***p*** |
| (Intercept) | 2.17 | 1.13 - 3.21 | **<0.001** |
| Day | 0.21 | 0.15 - 0.27 | **<0.001** |
| Type (co-axial) | 0.95 | 0.13 - 1.78 | **0.024** |
| Layers | -0.19 | -0.40 - 0.02 | 0.079 |
| **Random Effects** | | | |
| $\sigma^2$ | 0.40 | | |

| | |
|---|---|
| $\tau_{00\ ID}$ | 1.40 |
| $\tau_{11\ ID.day}$ | 0.02 |
| $r_{01\ ID}$ | -0.75 |
| ICC | 0.67 |
| N $_{ID}$ | 39 |
| Observations | 98 |
| Marginal $R^2$ / Conditional $R^2$ | 0.351 / 0.781 |

**Table S4.** Results from the random effects model in Table S3 with values transformed with an exponential function. This is necessary since the dependent variable is the logarithm of the force. Values calculated from the *lmer*() object from Table S3 and the *confint*() function in R (v. 4.3.1). Confidence intervals are calculated using profile-likelihoods for generalized linear models.

| Variable | Effect size | STD | t value | Effect size (exponential) | *p*-value | CI (inf.) | CI (sup.) | CI (inf., exponential) | CI (sup., exponential) |
|---|---|---|---|---|---|---|---|---|---|
| **Intercept** | 2.1712517 | 0.5226716 | 4.154141 | 8.769253 | 3.27E-05 | 1.163866 | 3.179135 | 3.202288 | 24.02597 |
| **Day** | 0.2138191 | 0.0305982 | 6.987971 | 1.238399 | 2.79E-12 | 0.152417 | 0.275496 | 1.164646 | 1.317184 |
| **Type(co-axial)** | 0.9538899 | 0.4168225 | 2.28848 | 2.595787 | 2.21E-02 | 0.127449 | 1.773505 | 1.135927 | 5.891469 |
| **Layers** | -0.1899968 | 0.1067994 | -1.77901 | 0.826962 | 7.52E-02 | -0.39763 | 0.018318 | 0.671911 | 1.018486 |

List of Primers used for RT-qPCR analysis:

**GAPDH:**
FW (5’ ATGGTGAAGGTCGGTGTGAA 3’)
RV (5’ GAGGTCAATGAAGGGGTCGT 3’)

**MyoD:**
FW (5’ CCACTCAGGTCTCAGGTGTAAC 3’)
RV (5’ TCGCCCGCTTGAGGAATAA 3’)

**Myogenin:**
FW (5’ CCCTACAGACGCCCACAATC 3’)
RV (5’ ACCCAGCCTGACAGACAATC 3’)

**MyHCI:**

FW (5’ GCCCCAAGCACAAGGAGT 3’)
RV (5’ AGCCCCAAGAAATAAGGACAG 3’)

**MyHCIIa:**
FW (5’ GCAGAGACCGAGAAGGAG 3’)
RV (5’ CTTTCAAGAGGGACACCATC 3’)

**MyHCIIb:**
FW (5’ GAAGGAGGGCATTGATTGG 3’)
RV (5’ TGAAGGAGGTGTCTGTCG 3’)

**MyHCIIx:**
FW (5’ GCGACAGACACCTCCTTCAAG 3’)
RV (5’ TCCAGCCAGCCAGCGATG 3’).
